# A prognostic signature for lung adenocarcinoma in people who have never smoked

**DOI:** 10.1101/2025.08.13.670169

**Authors:** Wei Zhao, Tongwu Zhang, Xing Hua, Phuc H. Hoang, Mona Miraftab, Monjoy Saha, John P. McElderry, Jian Sang, Olivia W. Lee, Caleb Hartman, Azhar Khandekar, Sunandini Sharma, Frank J. Colón-Matos, Samuel Anyaso-Samuel, Difei Wang, Kristine Jones, Amy Hutchinson, Belynda Hicks, Jennifer Rosenbaum, Xiaoming Zhong, Yang Yang, Angela Pesatori, Dario Consonni, David C. Christiani, Kin Chung Leung, Maria Pik Wong, Marta Manczuk, Jolanta Lissowska, Beata Świątkowska, Anush Mukeria, Oxana Shangina, David Zaridze, Ivana Holcatova, Dana Mates, Sasa Milosavljevic, Simona Ognjanovic, Milan Savic, Millica Kontic, Valerie Gaborieau, Paul Brennan, Oscar Arrieta, Yohan Bossé, Eric S. Edell, Matthew B. Schabath, Paul Hofman, Luis Mas, Sai S. Yendamuri, Chih-Yi Chen, I-Shou Chang, Chao Agnes Hsiung, Geoffrey Liu, Jacobo Martínez Santamaría, Bonnie E. Gould Rothberg, Karun Mutreja, Scott Lawrence, Nathaniel Rothman, Ludmil B. Alexandrov, Charles Leduc, Marina K. Baine, Philippe Joubert, Lynette M. Sholl, William D. Travis, Robert Homer, Qing Lan, Stephen J. Chanock, Lixing Yang, Soo-Ryum Yang, Jianxin Shi, Maria Teresa Landi

## Abstract

Knowledge of tumor cell dynamics can inform prognosis and treatment yet is largely lacking for lung adenocarcinoma in people who have never smoked (NS-LUAD). With RNA-seq data from 684 NS-LUAD and validation in an independent dataset, we identified three subtypes with distinct phenotypic traits and cell compositions. Additional genomic and histological data further characterized the subtypes. *‘Steady’*, marked by low proliferation, high alveolar cell fraction, moderate-to-well differentiation, and fewer driver genes’ alterations, is linked to prolonged survival and low immune evasion. *‘Proliferative*’ shows high proliferation markers, *TP53* mutations, and gene fusions. *‘Chaotic’*, with high epithelial-to-mesenchymal transition markers, has the worst prognosis even within stage I tumors. Lacking known molecular or histological characteristics, this aggressive subtype is solely identified by transcriptomic data. A 60-gene signature recapitulates the overall classification and strongly predicts survival even within subgroups based on tumor stage or known genomic features, emphasizing its potential for improving NS-LUAD prognostication in clinical settings.

**Significance:** The transcriptome of 684 lung adenocarcinomas in people who have never smoked (NS-LUAD) identifies three subtypes with different cellular dynamics, and genomic and morphologic features. A 60-gene signature accurately stratifies subjects for mortality risk, even in stage I, offering a clinically applicable tool for treatment decision making in NS-LUAD patients.

## Introduction

Approximately 10-25% of lung cancers occur in people who have never smoked(1,2), constituting one of the leading cause of cancer mortality worldwide(3). The majority of these are lung adenocarcinomas (LUAD)(4). Compared to smoking-related lung cancers, LUAD in people who have never smoked (NS-LUAD) is most common in women and the Asian population, and appears to differ in morphological presentations, molecular features and clinical outcomes(5,6). Many NS-LUAD cases have no identifiable risk factors and remain dramatically understudied.

In a previous analysis(5), we characterized the genomic alterations and evolutionary history of lung cancers in people who have never smoked using bulk whole-genome sequencing (WGS) data. We further expanded the analysis incorporating RNA-seq data, since accumulating evidence indicates that genomic changes alone may not be sufficient to drive tumor initiation and progression(7). In fact, somatic driver mutations have been identified in healthy tissues(8,9) and the presence of rare altered cell states has been identified in tumors with no clear genetic or somatic alterations(10,11). Acquisition of stem cell-like features by reactivating developmental programs contributes to metastasis, therapeutic resistance, and poor outcomes in lung cancer(12–15). Moreover, the tumor microenvironment (TME) is now widely appreciated as a pivotal player during tumor initiation, development, metastasis, and therapeutic resistance. A comprehensive investigation of the cell state plasticity and the complex cellular composition of the NS-LUAD and its TME through transcriptome profiling is critical for understanding tumor evolutionary patterns, improving the prediction of clinical outcomes, and informing treatment decision making.

Previous LUAD gene expression studies were predominantly conducted in patients’ samples with smoking history(16–21) and full transcriptomic sequencing (RNA-seq) was only examined in a few dozens of NS-LUAD per cohort, mostly of Asian ancestry (**Supplementary Table S1**). Here, we analyzed RNA-seq from 684 treatment-naïve NS-LUAD, including 386 (56.4%) stage I tumors, of both European and East Asian ancestry. With this unprecedented sample size, we identified three gene expression-based subtypes that encapsulate a combination of features, such as cell type composition, histological lineages, morphological characteristics, and genomic drivers. Importantly, we found substantial differences in survival and predicted treatment responses across these subtypes. Further, we developed a 60-gene signature that recapitulates the expression subtype classification and its prognostic capability, which could be easily used in clinical settings, even in the absence of driver gene or detailed morphological data. We validated our findings in an independent cohort of 110 NS-LUAD samples from Chen et al.(22).

## Results

### The Sherlock cohort

We assembled RNA-seq data from 684 tumor samples, including 610 from the Sherlock-*Lung* study and 74 from the TCGA-LUAD study, hereafter referred to as the Sherlock cohort (**Supplementary Table S2**). Paired lung adjacent normal samples were also profiled by RNA-seq from 491 subjects from the Sherlock-*Lung* study and 7 subjects from the TCGA-LUAD study. The patients’ clinicopathological characteristics are summarized in **Supplementary Table S3**. The cohort has a median age at diagnosis of 64 years and includes 129 (18.9%) male and 555 (81.1%) female patients, similarly between the Sherlock-*Lung* and TCGA-LUAD studies (**Supplementary Fig. S1A-B**). Forty subjects (5.8%) had stage IV disease at the time of diagnosis. In non-mucinous NS-LUAD, histologic grade based on the International Association for the Study of Lung Cancer (IASLC) grading system^32^ was evaluated in 301 cases with both RNA-seq data and H&E morphological images by a team of lung pathologists, resulting in 34 (11.3%) well, 69 (22.9%) moderate, and 198 (65.8%) poorly differentiated tumors (**Supplementary Table S2**).

### Gene expression pathways do not differ by ancestry/ethnicity

Using WGS-derived genetic ancestry, among the 610 subjects from Sherlock-*Lung*, 209 are of European ancestry, not Hispanic or Latino, from the US, Canada and Europe (referred to as EUR); 320 are of East Asian ancestry from East Asia, US and Canada (referred to as EAS); 15 are of native American/mixed ancestry from Europe and Canada (referred to as AMR or Mixed); and one is of African ancestry from the US (referred to as AFR). For the subjects with no WGS data, we used self-reported race/ethnicity and identified an additional 27 EUR, 20 EAS, 17 AMR or Mix and one patient of unknown ancestry. Among the 74 subjects from TCGA-LUAD, genetic ancestry was derived from genotyping data, but the geographical location is unknown. This cohort includes 59 EUR, two EAS, three AFR, and ten patients of unknown ancestry. In total, we examined 295 EUR, 342 EAS, 4 AFR, 32 AMR/Mix, and 11 with unknown ancestry or race/ethnicity (**Supplementary Fig. S1C**). Only 80 genes were differentially expressed between EUR and EAS tumors (the two groups with the largest sample size) and there was no significant difference in gene expression pathways between them (**Supplementary Fig. S2**). Therefore, we combined all ancestry/ethnic groups in further analyses.

### New transcriptomics-based classification identifies three major subtypes in NS-LUAD

We defined gene expression-based subtypes in NS-LUAD using non-negative matrix factorization (NMF), an approach that does not assume lack of correlation or independence and provides results with high interpretability(23). We assessed the stability of the decomposition results and found that a factorization rank of three and four (corresponding to three and four clusters) provide similar robust clustering (**Fig. 1, Supplementary Fig. S3, Methods**). As we aim to identify the minimum number of classes that could explain the inter-patient heterogeneity, we clustered the cohort into three groups.

**Fig. 1.**
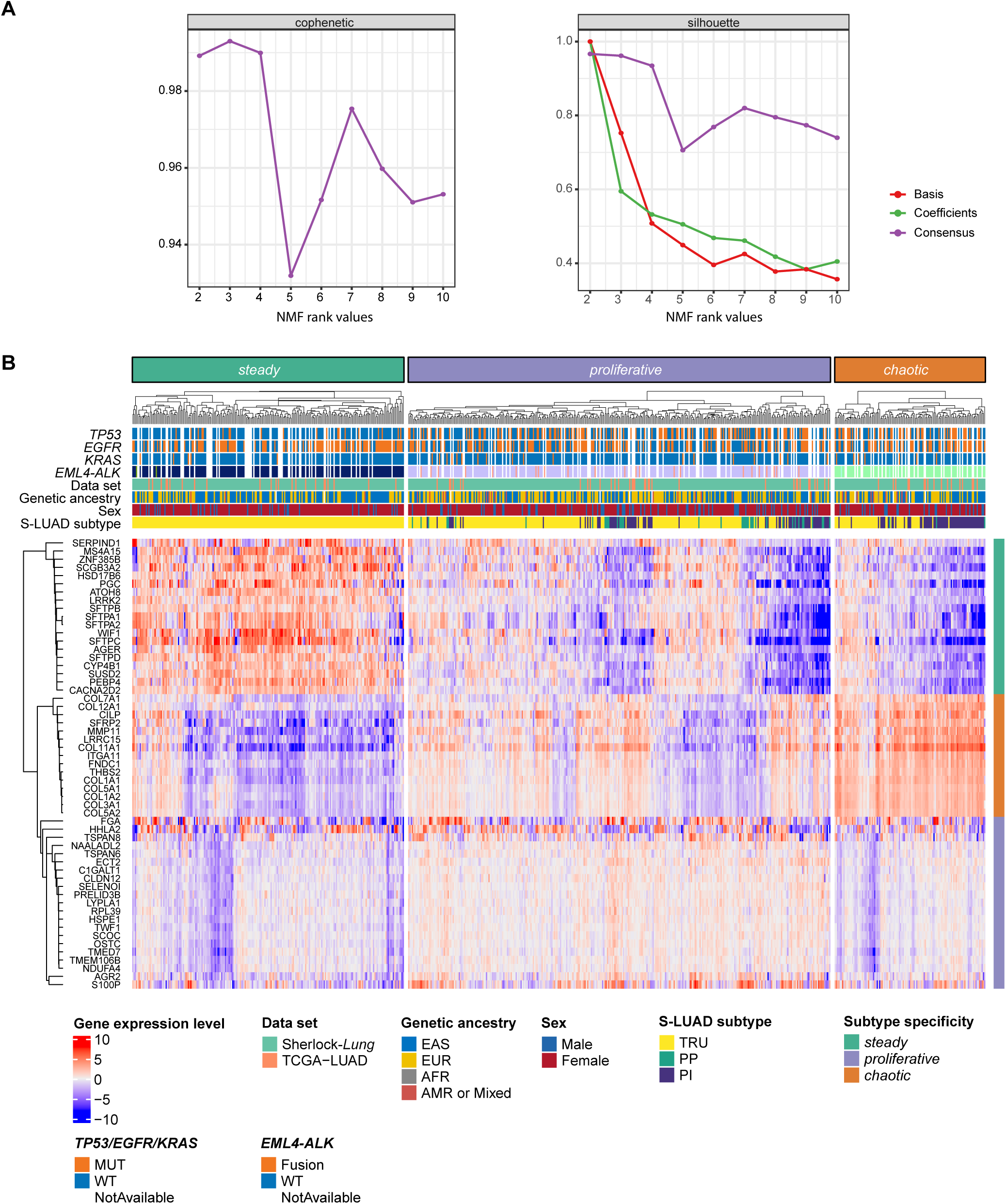
Transcriptomic clusters in the Sherlock cohort. (**A**) Consensus matrices for NMF rank 2 to 10. Cophenetic coefficient from consensus matrix (left) and silhouettes from basis, coefficient and consensus matrices (right) were computed from 80 runs for each rank value. (**B**) Clustering of subtype-specific genes in the three expression subtypes are defined by NMF. The top panel shows cluster assignment, driver mutations and fusions, data set, genetic ancestry, sex and S-LUAD subtypes. The right panel indicates the subtype specific genes.

The top genes contributing to the transcriptomics clustering include genes involved in lung development, cell-cell adhesion, epithelial-to-mesenchymal transition (EMT) pathways and immune responses (**Supplementary Table S4**). We performed pathway enrichment analysis in each subtype using the Ingenuity Pathway Analysis (IPA)(24) and Gene Set Enrichment Analysis (GSEA)(25) (**Supplementary Fig. S4-5, Supplementary Table S5, Methods**). The first cluster, named here ‘*steady’*, has a ‘normal-like’ expression pattern compared to other NS-LUAD tumors, with features similar to non-neoplastic tissues, including low expression levels of genes involved in proliferation, cell cycle and cell movement (**Supplementary Fig. S4A-B, S5A-B**), and low proliferation marker *MKI67* (**Supplementary Fig. S5C**). This subtype was also associated with significant upregulation of alveolar epithelial cell genes (**Supplementary Fig. S5D**). The second cluster, named ‘*proliferative*’, shows upregulation of proliferation signatures (e.g. genes regulated by *MYC*) and pathways involved in cell cycle and stemness features (**Supplementary Fig. S4C-D, 5E**). The third cluster, ‘*chaotic*’, shows high expression of genes associated with cell movement, EMT and cell adhesion (**Supplementary Fig. S4E-F, S5F-G**). The canonical mesenchymal markers *CDH2* and *TWIST1* were also highly expressed in *chaotic* (**Supplementary Fig. S5H-I**). To exclude the possibility that the gene expression differed purely as a consequence of different tumor purities, we repeated the analysis of the association of canonical proliferation and mesenchymal markers with the three clusters within tumors with estimated purity > 0.8 (**Methods**). The analysis showed consistent results and supported differentially regulated pathways in the NS-LUAD clusters (**Supplementary Fig. S6**).

Previous studies had defined three gene expression-based subtypes by unsupervised clustering of smoking-related LUAD (S-LUAD), including terminal respiratory unit (TRU), proximal-proliferative (PP), and proximal-inflammatory (PI)(26–28), with biological and clinical significance(17,29). We compared the three NS-LUAD expression subtypes to the previously defined S-LUAD expression subtypes (**Supplementary Fig. S7A**). We found that 100%, 67% and 46% of *steady*, *proliferative* and *chaotic* subtypes, respectively, were classified as TRU (**Supplementary Fig. S7B**), while in the TCGA S-LUAD(17), only 35% were TRU. Only 43 samples in this study were classified as PP and were mostly included in *proliferative* subtype. These results demonstrate that NMF clustering provides a more informative subtyping, and that NS-LUAD whole transcriptome substantially differ from S-LUAD.

### Clinical and morphological features of NS-LUAD transcriptomics-based subtypes

S*teady* tumors were enriched with stage I (74%, **Fig. 2A**, stage I proportions in *steady* vs. others: p=2.75x10^-8^), while 2/3 of *chaotic* had advanced stage (proportion of advanced stages vs stage I tumors in *chaotic* vs. other subtypes, p=1.44x10^-8^). Moreover, *steady* had significantly lower grade than the other tumors, with 53%, 27% and 23% of tumors being well/moderately differentiated in *steady*, *proliferative* and *chaotic* subtypes, respectively (**Fig. 2B**, *steady* vs others, p=5.52x10^-6^).

**Fig. 2.**
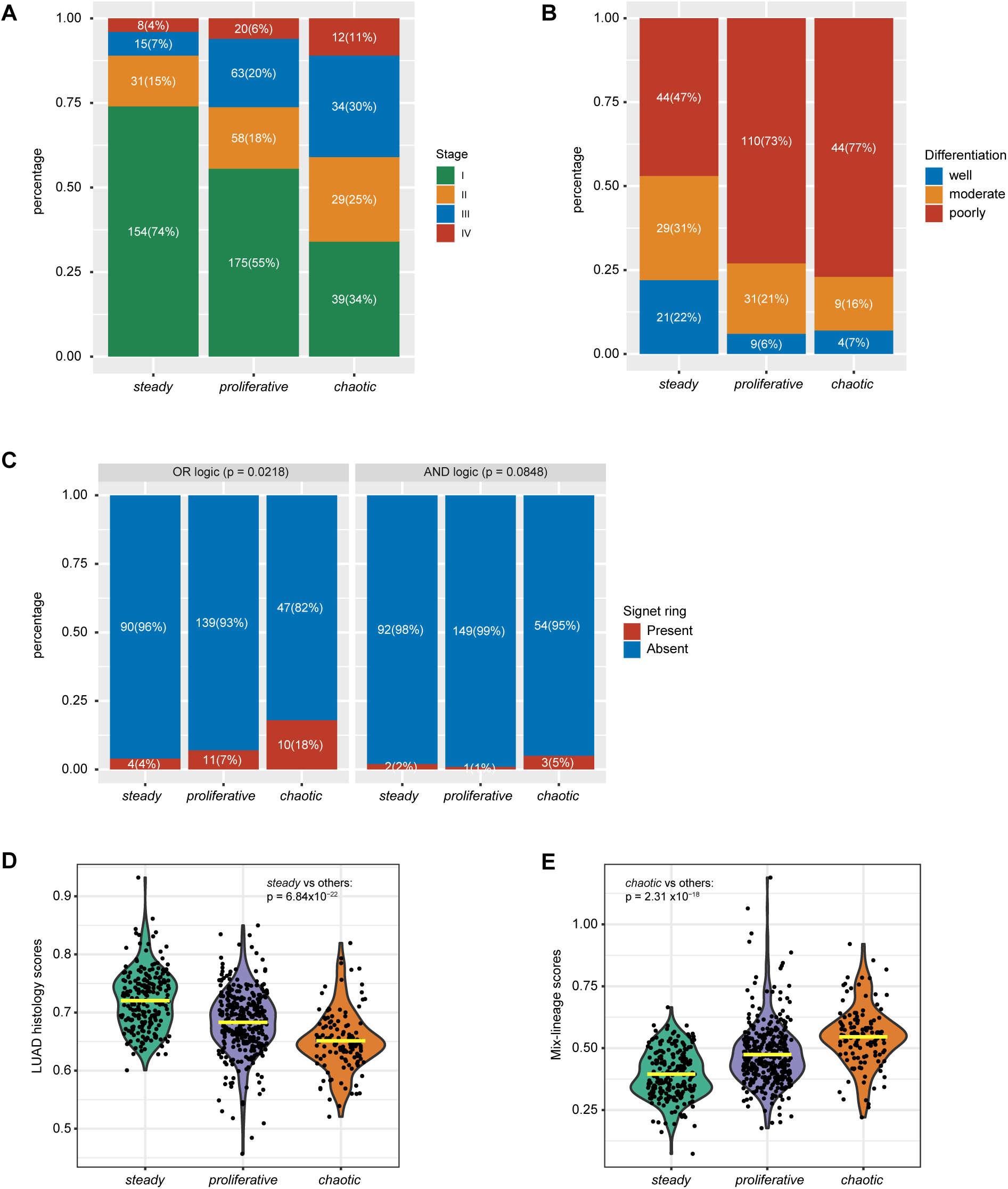
Comparison of clinical and histopathological features across NS-LUAD expression subtypes. **A-C**, Comparison of (**A**) tumor stages, (**B**) histologic grades, and (**C**) signet ring cell features across the NS-LUAD expression subtypes. P-values for comparison of morphological features from chi-squared test are shown. **D-E**, Violin plots of (**D**) LUAD histology scores and (**E**) mix-histological lineage scores across the NS-LUAD expression subtype. Mean values are indicated by the yellow lines. P-values from two-sided Mann–Whitney U-test are shown.

We assessed the presence of six histologic/cytologic features in the three subtypes, including lymphoid aggregate, tumor spread through air spaces, and clear cell, morular, mucinous, and signet ring cell (SRC) features (**Methods**). In line with the previous findings that the SRC feature in lung cancer is associated with high histologic grade and worse survival(30–32), we observed that the *chaotic* tumors had higher fractions of SRC^+^ tumors (**Fig. 2C**); they also tended to lack clear cell features (**Supplementary Fig. S8**).

### Cell lineage infidelity and cellular plasticity increases from *steady* to *chaotic* subtypes

Disruption of lineage integrity has been identified as an important driver of lung cancer initiation and progression(13,15). We analyzed the compositions of cell lineages typical of three histological types - LUAD, lung squamous cell carcinoma (LUSC) and neuroendocrine tumor (NET) - in each tumor using lineage-specific genes defined by single-cell RNA-seq (scRNA-seq) (**Supplementary Table S6**)(33). We assessed the presence of mix-lineage based on scores from the three histological types, with high scores representing high lineage-mixing features. While all the tumors had higher scores for LUAD compared to LUSC or NET, as expected, *steady* had significantly higher LUAD scores compared to the other subtypes (**Fig. 2D**, p=6.84x10^-22^), indicating a high fidelity to the LUAD lineage. In contrast, *chaotic* had higher mixed-lineage scores, even though histologically they were LUAD (**Fig. 2E**, p=2.31x10^-18^). Interestingly, while the LUAD lineage component slightly decreased with stage in general, among the three subtypes the decreasing trend was observed only in *chaotic* (**Supplementary Fig. S9**).

Overall, *steady* was associated with lower tumor stage and histologic grade, and exhibited higher histological lineage fidelity, while *chaotic* was associated with advanced tumor stage, signet ring cells, and mixed lineage.

### The proportion of epithelial cells and cancer-associated fibroblasts differ across subtypes

Previous studies have identified substantial tissue heterogeneity in lung cancer(15,34–41). Here we estimated the abundances of major cell types by cell deconvolution using bulk RNA-seq data and expression signature genes from a LUAD scRNA-seq study(41) (**Methods**). *Proliferative* and *chaotic* had higher proportions of epithelial cells (**Fig. 3A-B**, mean=38.33%, p=2.03x10^-12^ in *proliferative* vs. others), and fibroblasts (**Fig. 3C**, mean=21.55%, p=6.88x10^-40^ in *chaotic* vs. others), respectively. An inverse association between fibroblasts and morphological lepidic components was present as expected (**Supplementary Fig. S10**). The association of *chaotic* with fibroblasts remained statistically significant after accounting for tumor lepidic features (p=2.57x10^-19^, **Methods**)

**Fig. 3.**
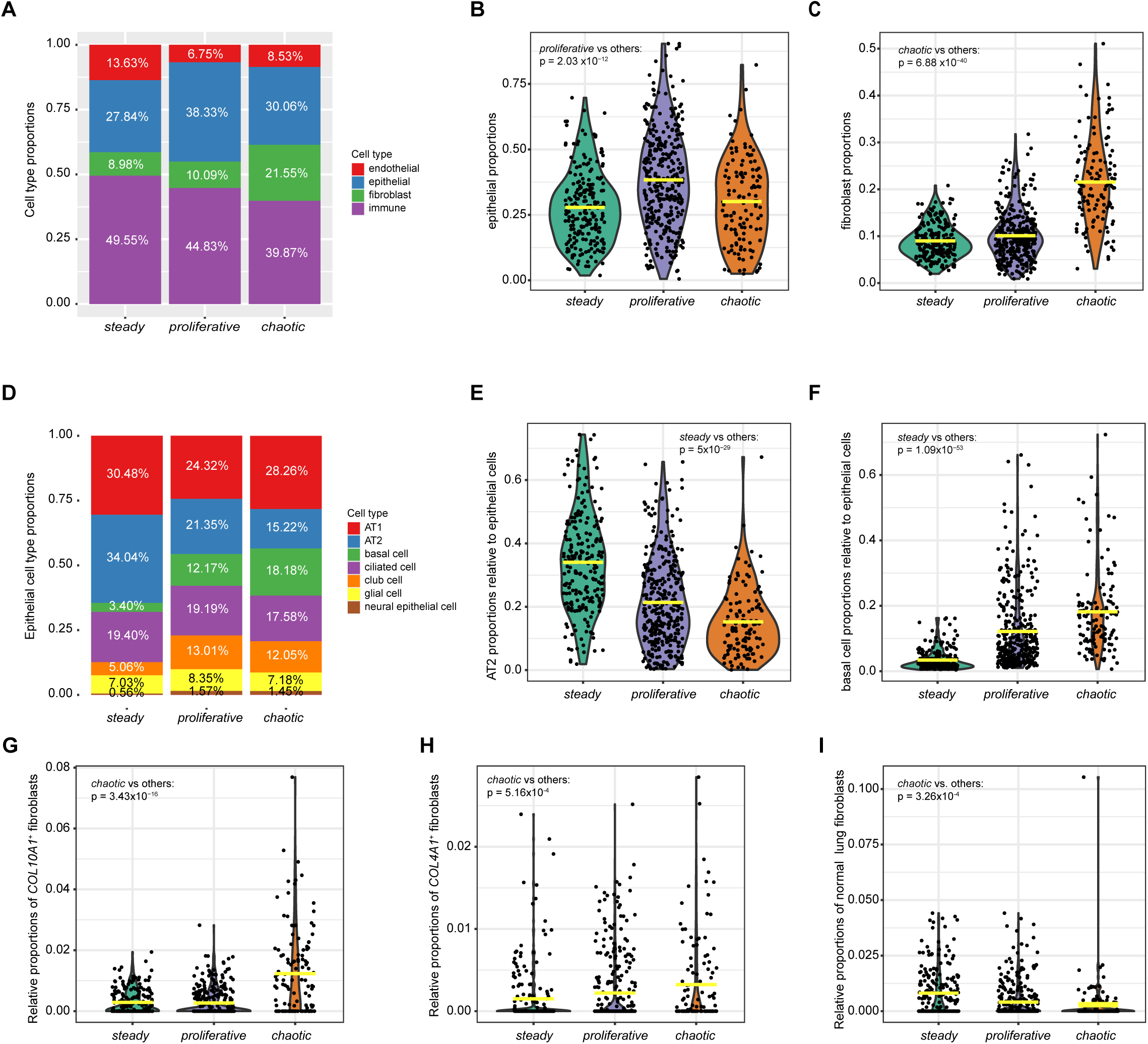
Cell type deconvolution in the Sherlock cohort. **A**, Summary of the average proportions of cell types in NS-LUAD expression subtypes. **B-C**, Violin plots of the proportions of (**B**) epithelial cells and (**C**) fibroblasts across NS-LUAD expression subtypes. **D**, Summary of the average proportions of epithelial cell types relative to all epithelial cells. **E-F**, Violin plots of the proportions of (**E**) AT2 and (**F**) basal cells relative to all epithelial cells across NS-LUAD expression subtypes. **G-I**, Comparison of the relative proportions of (**G**) *COL10A1*+, (**H**) *COL4A1*+, and (**I**) non-malignant lung fibroblasts across NS-LUAD expression subtypes. The fibroblast estimates are relative differences in abundances and cannot be compared across cell types. Mean values are indicated by the yellow lines in the violin plots. P-values from two-sided Mann-Whitney U-test are shown.

Among epithelial cells, *steady* had substantially higher proportions of alveolar AT2 cells (**Fig. 3D-E**, mean=34.94%, p=5x10^-29^ in *steady* vs. others), consistent with the observed upregulation of the alveolar cell signature in *steady* (**Supplementary Fig. S5D**) and the high score for LUAD lineage (**Fig. 2D**).

Correspondingly, *steady* was depleted of basal cells (**Fig. 3F**, mean=3.40%, p=1.09x10^-53^ in *steady* vs. others). Among the alveolar cell types, the high levels of *SFTPB*, *SFTPC* and other markers(41) shows preponderance of AT2 features. For simplicity, we define these cells as AT2 throughout the manuscript; however, some AT1 cells undergoing reprogramming towards AT2 cells may be included in this category.

To dissect the fibroblast cell types in *chaotic*, we decomposed the data based on a scRNA-seq study of stromal cells in lung cancer(36) using a computational algorithm(42) that allows estimation of relative cell abundances with high accuracy (**Fig. 3G-I**, **Methods**). Compared to the other subtypes, we found that *chaotic* had significantly higher relative abundances of two classes of cancer-associated fibroblasts (CAFs), including the fibroblasts expressing *COL10A1* (*COL10A1*^+^) and fibroblasts expressing *COL4A1* (*COL4A1*^+^), and was depleted of non-malignant fibroblasts. Notably, given the different stage distribution across subtypes, we verified that the association of cell compositions with subtypes remained significant within each stage (**Supplementary Fig. S11**). The expression of canonical cell type markers confirmed the cell composition across the three subtypes (**Supplementary Fig. S12**).

### Driver genes and fusions strongly differ across transcriptomics subtypes

To verify whether the gene expression-based subtypes harbored mutational driver events and to explore possible therapeutic opportunities in NS-LUAD subtypes, we integrated RNA-seq data with WGS data from 499 tumors, including 486 tumor samples from the Sherlock-*Lung* study and 13 from the TCGA-LUAD (**Fig. 1, Supplementary Fig. S13-S14**). *TP53* mutations were significantly enriched in *proliferative* (p=4.99x10^-6^, FDR=9.73x10^-4^), while were highly depleted in *steady* (p=4.63x10^-14^, FDR=9.03x10^-12^). Whole genome doubling (WGD), often found in association with *TP53* mutations across cancers(5), was not enriched in *proliferative*. *ALK* fusions, often associated with a tumor aggressive behavior(43), were negatively associated with *steady* (p=1.73x10^-4^, FDR=0.0168). Co-occurrence of *ALK* fusions and *TP53* mutations, known to be associated with poor prognosis and treatment resistance(44,45), was observed in seven *proliferative* tumors. Although not significantly after multiple testing correction, *EGFR* mutations were more frequent in *steady* (p=3.65x10^-3^, FDR=0.315), consistent with our previous observation of a slow tumor growth in EGFR-positive tumors(5). No driver mutations or fusions (e.g., *MET* exon 14 skipping) were significantly enriched in *chaotic*, although *TP53* mutations and WGD were frequently found.

We further investigated gene fusions beyond driver events, most of which were validated with WGS data (**Supplementary Table S7**). A total of 11,947 fusion events were identified using RNA-seq in 638 out of 684 tumors, including 64.6% intrachromosomal and 35.4% interchromosomal fusions (**Supplementary Fig. S15A, Supplementary Table S7**). Substantially more fusion events were detected per sample in *proliferative* (mean=22.1) compared to *steady* (mean=13.9) and *chaotic* (mean=10.7) tumors (p=1.11 x 10^-13^, **Fig. 4A**). In tumors with available WGS data, 54.3% of fusions were supported by structural variants detected by WGS, similarly across the three subtypes (**Supplementary Fig. S15B**). Among the fusions, 3,422 (28.6%) were detected in pairs of protein-coding genes and 953 (8.0%) were in-frame (**Fig. 4B, Supplementary Fig. S15C**). We detected 39 recurrent protein-coding fusion events (defined as fusions between pairs of protein coding genes detected in more than one tumor, **Fig. 4C**). The most frequently occurring in-frame fusions were *EML4*-*ALK* (33 tumors) and *KIF5B*-*RET* (8 tumors). The *PARG-BMS1* fusion identified in five tumors is a novel recurrent fusion event in lung cancer. It involves *PARG*, whose protein product is important for DNA damage repair(46), and has been previously reported in breast cancer(47). Consistent with previous studies(17,22), genes that were most frequently involved in in-frame fusions include *ALK* (35 tumors), *EML4* (33 tumors), *RET* (12 tumors) and *ROS1* (9 tumors) (**Fig. 4D**). Tumors in the *proliferative* subtype harbored significantly more protein-coding and in-frame fusions per sample than the other subtypes (**Supplementary Fig. S15D-E**), and among them, a higher frequency of in-frame *ALK* fusions (8.2%) compared to *steady* (2.3%) and *chaotic* (1.6%) (**Fig. 4E**), showing consistency between RNA-seq and WGS-based results.

**Fig. 4.**
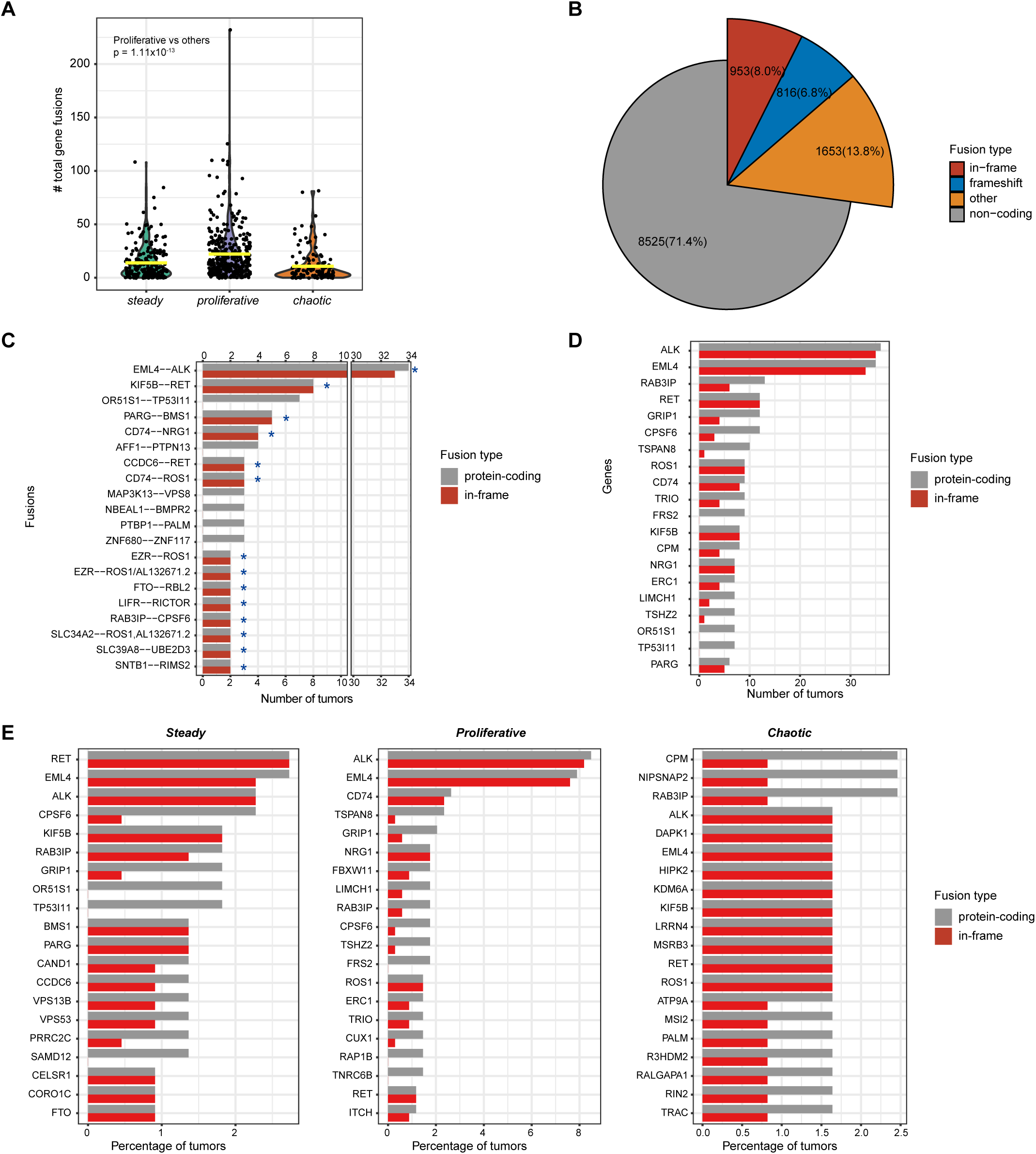
Gene fusion events in the Sherlock cohort. (**A**) Comparison of the total numbers of gene fusions per sample across NS-LUAD expression subtypes. Mean values are indicated by the yellow lines. P-value from two-sided Mann-Whitney U-test is shown. (**B**) Summary of types of all detected gene fusions. (**C**) Summary of frequent gene fusions between pairs of protein-coding genes. The bar plot indicates the number of tumors containing gene fusion events. Colors indicate fusion types. Recurrent in-frame fusions of protein-coding genes are indicated by the star signs(*). (**D**) Summary of genes most frequently involved in fusions between protein-coding genes. The bar plot indicates the number of tumors in which gene fusions involving the listed genes are detected. Colors indicate fusion types. (**E**) Summary of genes most frequently involved in fusions between protein-coding genes within NS-LUAD expression subtypes. The bar plots indicate the percentage of tumors in which gene fusions involving the listed genes are detected. Colors indicate fusion types.

### Dramatic survival differences across subtypes even within stage I

To evaluate the prognostic power of the NS-LUAD expression subtypes beyond established prognostic factors, we tested mortality risk using multivariable Cox models adjusting for age, sex and tumor stage in all Sherlock subjects and within subgroups defined by molecular and clinical classifiers (**Fig. 5**). We analyzed the 10-year overall survival rates for all subtypes and further analyzed the 5-year overall survival rates for *chaotic* due to its association with advanced stage. We found that *steady* had better survival (HR=0.43, 95% CI=0.3-0.63, p=1.3x10^-5^), while *chaotic* had worst survival (HR_10-year_ =1.4, 95% CI=0.99-2, p=0.056; HR_5-year_ =1.5, 95% CI=1.0-2.3, p=0.031). Given the high prevalence of early and late tumor stages within *steady* and *chaotic*, respectively, we restricted the analyses to stage I tumors only (n=374) and confirmed the subtype associations with survival (*Steady*: HR=0.39, 95% CI=0.22-0.68, p 8.8x10^-4^; *chaotic*: HR_10-year_ =2.5, 95% CI=1.3-4.7, p=0.0043; HR_5-year_ =3.7, 95% CI=1.8-7.3, p=2.2x10^-4^), indicating that the transcriptomic profiling captures subgroups of stage I tumors that are more aggressive and may benefit from *ad hoc* treatments.

**Fig. 5.**
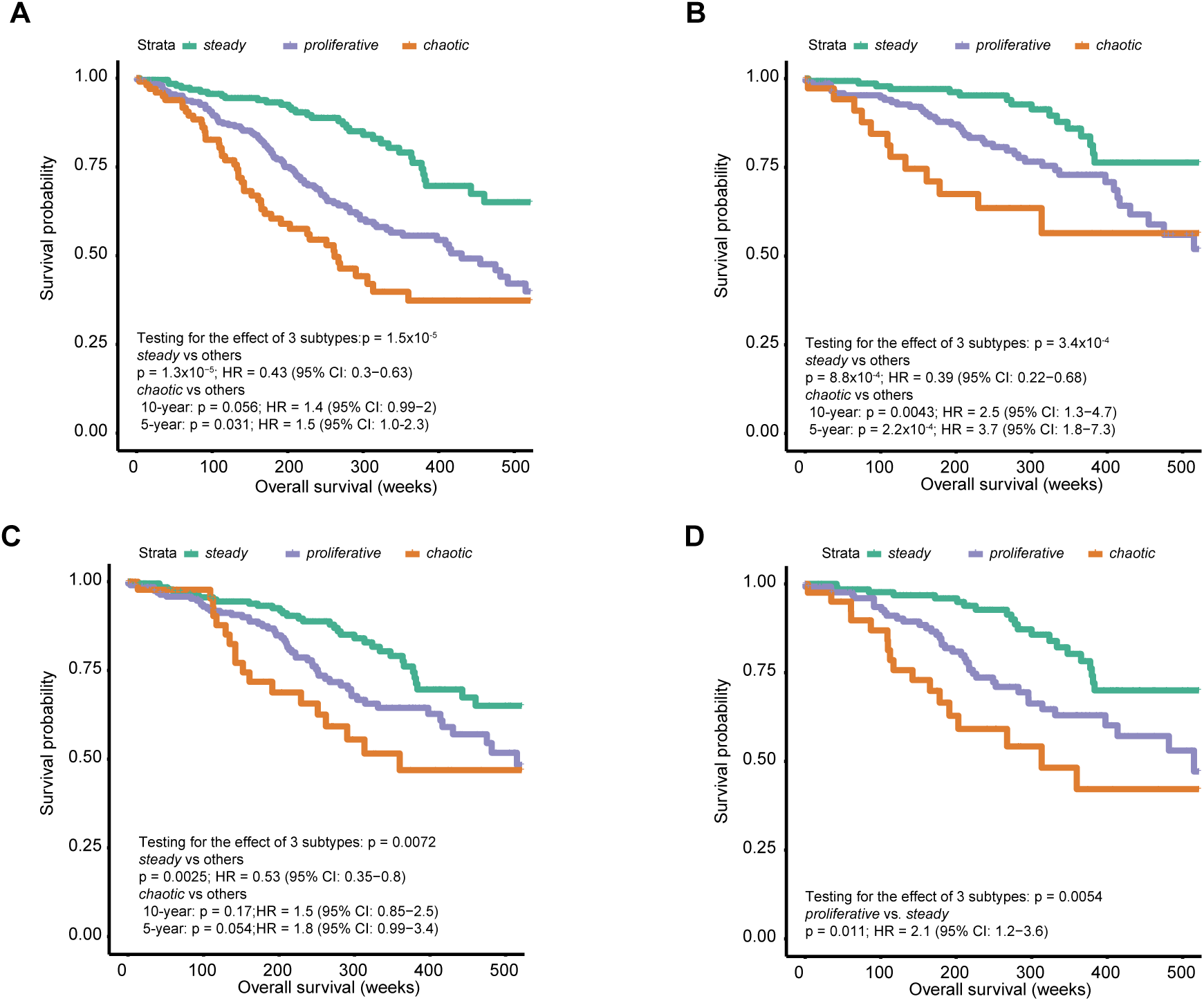
Association between NS-LUAD expression subtypes and overall survival in the Sherlock cohort. **A-D**, Kaplan-Meier survival curves for overall survival stratified by NS-LUAD expression subtypes in (**A**) all patients, (**B**) stage I patients, (**C**) patients classified as TRU subtype^26^, and (**D**) *TP53*-wt patients. P-values and hazard ratios (HR) were calculated using Cox proportional hazards models adjusting for age, sex, and tumor stage (the analysis restricted to tumor stage I did not include tumor stage in the model).

To test whether the NS-LUAD subtypes adds significant prognostic information to the previously defined S-LUAD subtypes, we restricted the analyses to the 303 tumors that were classified as TRU based on the S-LUAD classification. Even within the TRU tumor group, we could clearly see consistent prognostic differences across transcriptomic-based subtypes although the statistical significance was borderline for *chaotic* likely due to the small sample size (n =56) (*Steady*: HR=0.53, 95% CI=0.35-0.8, p=0.0025; *chaotic*: HR_10-year_ =1.5, 95% CI=0.85-2.5, p=0.17; HR_5-year_ =1.8, 95% CI=0.99-3.4, p=0.054). Moreover, clinical studies have suggested that non-small cell lung cancers with *TP53* alterations have worse prognosis(48,49). As the *TP53* mutations were enriched in *proliferative* and depleted in *steady*, we analyzed 169 tumors with no *TP53* mutations to verify whether the distinction in survival between *proliferative* and *steady* was solely driven by the *TP53* mutations. We found that tumors in *proliferative* were still associated with shorter survival than *steady* (HR=2.1, 95% CI=1.2-3.6, p=0.011).

### Predicted response to immune checkpoint blockade in the *steady* subtype

Compared to LUAD from patients with smoking history, NS-LUAD are thought to be less sensitive to immune checkpoint blockade (ICB) due to differences in tumor mutational burden and immune microenvironment, including *PD-L1* expression(50) – two established markers predictive of ICB responses(51–56). We tested whether transcriptional classes could identify NS-LUAD patients that could potentially benefit from ICB therapy. We predicted the responses to ICB using the Tumor Immune Dysfunction and Exclusion (TIDE) algorithm(57) in 498 tumors with paired tumor and normal RNA-seq data. A lower TIDE score predicts a better ICB response and has been shown to be more accurate than *PD-L1* levels or mutation load(57). Surprisingly, *steady* had lower TIDE scores than the other subtypes (**Supplementary Fig. S16A**, p=6.21x10^-14^). Compared to the paired normal samples, most NS-LUAD tumors (77.1%) had low levels of cytotoxic T lymphocytes (CTL) across all three NS-LUAD subtypes (**Supplementary Fig. S16B**). However, *steady* had lower abundances of suppressive cells prohibiting T cell infiltration, including myeloid-derived suppressor cells (MDSCs, p=4.77x10^-19^) and cancer-associated fibroblasts (CAFs, p=1.44x10^-10^), which are known mechanisms for tumor immune evasion through T cell exclusion(58,59) (**Supplementary Fig. S16C-D**). These findings are in line with the TIDE score results but warrant further validation.

Since ICB response rate is approximately 20% in unselected non-small cell lung cancer patients(60), we focused on the bottom 20% of the TIDE scores. *Steady* was enriched (**Supplementary Fig. S16E-F**, p=1.09x10^-9^) and *chaotic* was depleted (p=3.58x10^-4^) in this lowest TIDE quintile. We also measured the expression of *PD-L1*(*CD274*) and *PD-1*(*PDCD1*). As expected, the ICB prediction based on the TIDE scores was inversely correlated with *PD-L1*/*PD-1* expression. However, *PD-L1*/*PD-1* expression explained only a small proportion of the TIDE score variance (R^2^*_PD-L1_*=0.081, proportion of the variance explained by *PD-L1*, p*_PD-L1_*=1.06x10^-10^; R^2^*_PD-1_*=0.037, p*_PD-1_*=1.46x10^-5^; R^2^*_PD-L1+PD-1_*=0.093, p*_PD-L1+PD-1_*=3.87x10^-12^; **Supplementary Fig. S16E-F**). Adding the information on transcriptomics-based subtypes improved the ICB response prediction beyond that based on *PD-L1*/*PD-1* expression (R^2^_subtype+PD-L1+PD-1_=0.25, p=1.17x10^-22^, Supplementary Fig. S16G).

### A 60-gene signature of NS-LUAD subtypes with prognostic significance

To identify the minimum subset of genes capable of distinguishing the three subtypes for potential clinical application, we applied a centroid-based prediction method in the Sherlock cohort (**Supplementary Fig. S17, Methods**). To assess the gene set accuracy for predicting subtypes, we adopted a cross-validation approach within the Sherlock cohort and estimated the error rate as 13.2% and 15.1% in the training and testing sets, respectively (**Methods**). We obtained a 60-gene signature including approximately 30 genes involved in lung development and about 10 genes in EMT or cell-cell adhesion (**Supplementary Table S8**).

The 60-gene signature correctly predicted expression subtypes of 595 (87.0%) samples consistently with the NMF classification results based on the whole transcriptome (**Supplementary Table S9**). The tumors predicted as *steady* were significantly associated with prolonged overall survival after adjusting for age, sex and tumor stage, and remained significant when restricting the analysis to patients with stage I, TRU or *TP53*-wt tumors (**Supplementary Fig. S18**). The association between predicted *chaotic* tumors and worst survival was also confirmed in all tumors and within subgroups of stage I and *TP53*-wt patients.

We validated the performance of the 60-gene signature in an independent data set from the Genomic Institute of Singapore (referred to as the GIS cohort)(22) (**Supplementary Table S10**). Although a much smaller sample size (n=110 samples), the GIS cohort was the largest publicly accessible RNA-seq dataset from lung cancer in people who have never smoked and was limited to the Asian population. In the GIS cohort, the 60-gene signature classified 35, 64 and 11 tumors as *steady*, *proliferative*, and *chaotic*, respectively. Excluding *chaotic* because of the small sample size, the 60-gene signature confirmed mortality risk prediction after adjusting for age, sex and tumor stage (c-index=0.682), with *steady* significantly associated with prolonged survival overall (**Fig. 6A**, HR=0.32, 95% CI=0.12-0.87, p=0.026) and within *TP53*-wt tumors (**Fig. 6B**).

**Fig. 6.**
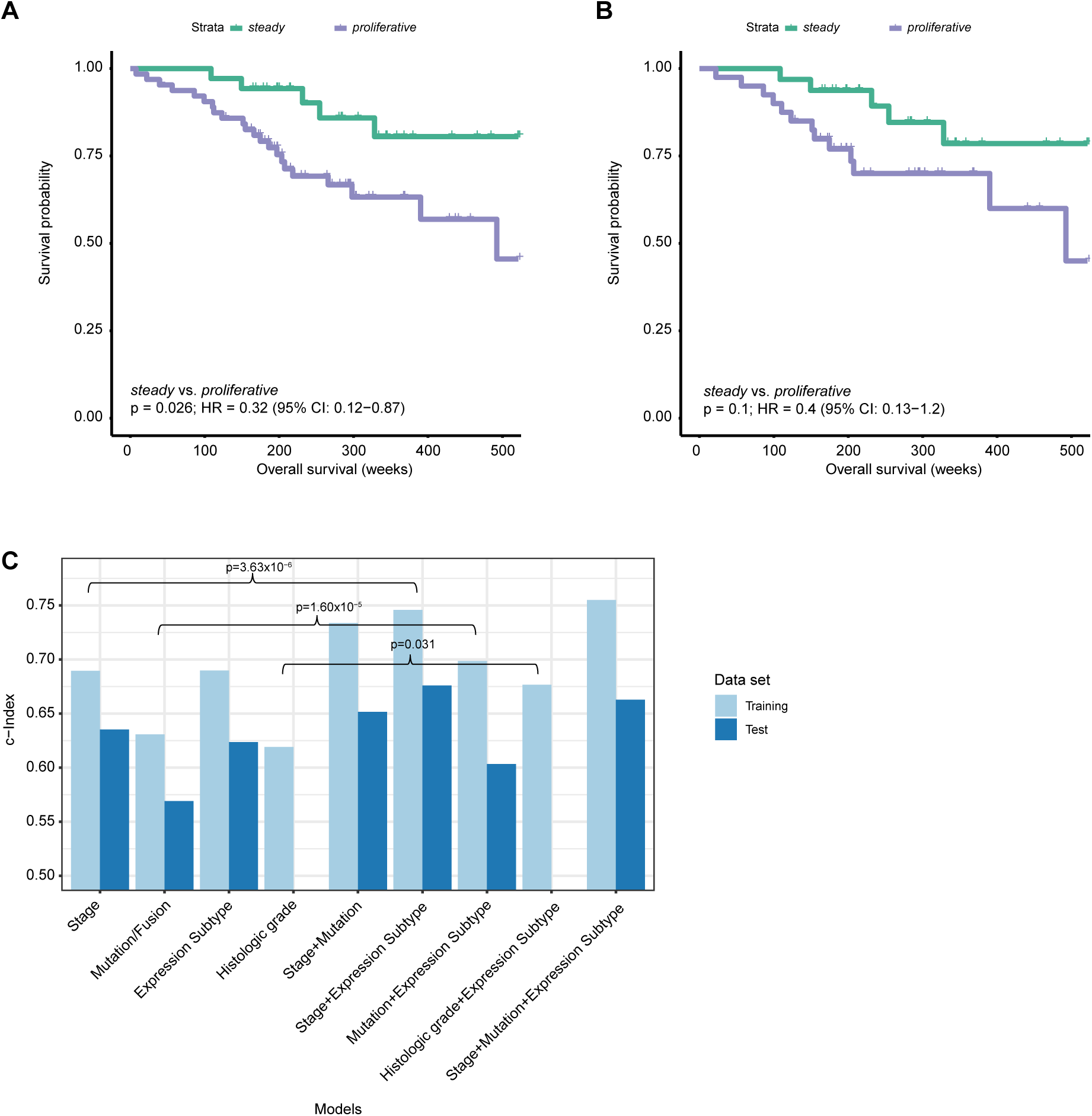
Association between NS-LUAD expression subtypes and overall survival in the Genome Institute of Singapore (GIS) cohort^22^. **A-B**, Kaplan-Meier survival curves for overall survival stratified by NS-LUAD expression subtypes in (**A**) all patients and (**B**) *TP53*-wt patients. P-values and hazard ratios (HR) are calculated using Cox proportional hazards models adjusting for age, sex, and tumor stage. (**C**) C-indexes of Cox models predicting the overall survival using different sets of predictors in the training (Sherlock) and testing (GIS) data sets. P-values testing for the effect of adding the expression subtypes to models of single modality were calculated using likelihood ratio test.

To develop an overall prognostic risk score, we trained Cox regression models in the Sherlock cohort using (A) tumor stage (‘*Stage model’*), (B) driver mutations in *TP53*, *EGFR* or *KRAS* and *EML4*-*ALK* fusions (‘*Mutation model*’), (C) NS-LUAD expression subtypes quantified as the distance to the centroids of the three expression subtypes (‘*Expression Subtype model*’), (D) histologic grade (‘*Histologic grade* model’), (E) tumor stage plus mutations/fusions, (F) expression subtype plus stage, (G) expression subtype plus mutations/fusions, (H) expression subtype plus histologic grade, (I) expression subtype plus both stage and mutations/fusions. Each model was tested in the GIS set (**Fig. 6C**). Since the GIS cohort lacks morphological data, the predictive power of histologic grade compared to other models was assessed based on the performance in the training set. We found that among models of single modality (model A-D), the *stage* and *expression subtype* models had similar performance and were superior to the *mutation* and *histologic grade* models. Combining more features improved the predictive performance compared to single modality models. Notably, adding transcriptomics-based subtypes significantly improved the predictive power of the models based on *stage*, *mutation*, or *histologic grade,* respectively (**Fig. 6C**). Across all models, the one combining expression subtype plus stage provided the most accurate prediction of survival in the testing set (c-index=0.68).

## Discussion

Understanding tumor cell plasticity and composition and its TME can offer important insights into evolutionary trajectories, prognosis and treatment options. In this study, whole transcriptomic analysis of 684 lung adenocarcinomas from subjects who have never smoked identified three subtypes that encompass tumors’ genomic, clinical, and morphological features. The *steady* subtype is characterized by low proliferation markers, high proportion of alveolar epithelial cells, low histologic grade and depletion of *TP53* mutations and *ALK* fusions. The *proliferative* subtype, on the contrary, features upregulation of proliferation and cell cycle pathways and enrichment of *TP53* mutations, and *ALK* and other protein-coding gene fusions. The *chaotic* subtype is characterized by elevated levels of mesenchymal cell state, disrupted cell-cell adhesion, increased proportions of cancer associated fibroblasts and histologic features of mixed-lineage, but no significant enrichment of driver mutations or fusions, suggesting that the transcriptomic data capture features beyond the known drivers. Notably, this NMF clustering provides a more informative subtyping than the original TCGA transcriptional classification based on 500 genes and highlights large differences between NS-LUAD and from LUAD in subjects with smoking history. These subtypes have superb prognostic capability even within stage 1, identifying early-stage tumors with otherwise unrecognizable aggressive behavior that may require *ad hoc* treatment.

Previous studies mostly on subjects with smoking history identified gene expression signatures for LUAD prognosis (mostly based on a handful of genes and AI-approaches) and developed models to stratify LUAD patients into different prognostic groups(61–65). Here, the NS-LUAD transcriptomics-based subtypes show robust prognostic performance also within subgroups defined by tumor stage, driver gene mutations, and S-LUAD transcriptional classification, with *steady* associated with prolonged survival and *chaotic* with the poorest survival. Remarkably, including the information on the three subtypes into prognostic predictive models strongly improved survival prediction compared to the models using solely stage, grade differentiation or driver gene mutations. The combination of stage information with the expression subtypes provided the best prognostic predictive power. Furthermore, the subtype information significantly improves the prediction of ICB response beyond that based on *PD-L1*/*PD-1* expression, identifying a subgroup in *steady* with strong predicted response to ICB therapy. This indicates that the transcriptomics-based subtyping could be a tool for the ICB response prediction in NS-LUAD but needs validation in studies with both transcriptomic and ICB response data.

Finally, we demonstrated that the expression of just 60 genes can provide information on the underlining NS-LUAD architecture, encompassing histological, genomic and cell composition features, and to predict survival even within subgroups based on tumor stage or known molecular features. All major findings were validated in an independent cohort exclusively of east Asian ancestry. Validation in larger date sets representative of more heterogeneous genetic ancestries is warranted in future studies. Importantly, like the whole transcriptomics-based subtype classification, the 60-gene signature can discriminate aggressive from more stable tumors even within stage I, providing an easily adoptable tool for treatment decision making in hospital settings.

## Methods

### Ethics declarations

The NCI exclusively received de-identified samples and data from collaborating centers, had no direct interaction with study subjects, and did not use or generate any identifiable private information, therefore the Sherlock was classified as “Not Human Subject Research (NHSR)” according to the Federal Common Rule (45 CFR 46; https://www.eCFR.gov).

Some tissue specimens were obtained from the IUCPQ Tissue Bank, site of the Quebec Respiratory Health Network Biobank or the FQRS (https://www.tissuebank.ca) in compliance with Institutional Review Board-approved management modalities. All participants provided written, informed consent.

Some samples and data from patients included in this study were provided by the INCLIVA Biobank (PT17/0015/0049), integrated in the Spanish National Biobanks Network and in the Valencian Biobanking Network, and they were processed following standard operating procedures with the appropriate approval of the Ethics and Scientific Committees.

### Sample collection, RNA sequencing, and public data preprocessing

We collected 620 fresh-frozen tumor samples from 610 treatment-naïve lung cancer patients. Among these patients, matched fresh-frozen normal lung tissue samples were also obtained from 491 subjects. All tumors were reviewed by seven lung pathologists using FFPE tissue blocks to determine the histological subtypes and morphological features and were confirmed to be lung adenocarcinomas. RNA-seq was performed using the Illumina NovaSeq6000 platform (RRID:SCR_016387) and Roche KAPA RNA HyperPrep with RiboErase protocol, generating 2x151bp paired-end reads.

The FASTQ files were aligned to the human reference genome GRCh38/hg38 using STAR v2.7.3(RRID:SCR_004463)(66), and were quantified using HTSeq v2.0.4(RRID:SCR_005514)(67) and GENCODE v35 (RRID:SCR_014966) (68). Quality control was performed at three levels: (1) at FASTQ file level, the raw data was analyzed by FastQC (RRID:SCR_014583) (69). We assessed five main QC metrics (base-wise quality, k-mer overrepresentation, guanine-cytosine content, content of N bases and sequence quality) defined by the PCAWG study(18) and excluded samples if three or more metrics failed. (2) At alignment level, we assessed the BAM files using STAR and PicardTools(RRID:SCR_006525) (70) and excluded samples if more than 50% PF base pairs (base pairs that passed Illumina quality filtering) were unmapped or the total reads were fewer than 1 million. (3) At gene quantification level, we excluded samples with fewer than 5 million total counts. For each subject with multiple tumor samples, the sample with the largest number of PF reads (reads that passed Illumina quality filtering) was included in the analysis. The resulting expression data is based on 610 tumor samples and 491 matched normal samples.

For the TCGA-LUAD data set, raw RNA-seq FASTQ files were downloaded from the GDC legacy archive (RRID:SCR_014514) (71). 74 tumor samples and 7 matched normal samples from NS-LUAD subjects passed QC and were included in the analysis. For the GIS data set, raw RNA-seq FASTQ files from 110 tumor samples and 59 matched normal samples from 110 NS-LUAD subjects were downloaded from the European Genome-Phenome Archive (EGA, RRID:SCR_004944, http://www.ebi.ac.uk/ega/) under the accession code EGAD00001004421. All data sets were processed using the same pipeline for alignment, quantification and QC used for the Sherlock cohort.

The expression read counts from the three sets were processed by the ComBat-Seq (RRID:SCR_010974) (72) for batch adjustment followed by TMM normalization using DESeq2(73).

### Expression subtype detection

TMM normalized data with median gene levels greater than 1 was sorted by variance. The top 5000 genes with large variance excluding the mitochondrial genes were used for expression subtype detection. Subtypes were identified using the R package NMF (RID:SCR_023124) (74) and the Brunet’s algorithm(75). To estimate the factorization rank (equals to the number of clusters in this analysis) and to assess the stability of the results, we performed 80 runs of NMF analysis for the rank value 2 to 10.

The rank of 3 corresponding to three clusters was selected based on the cophenetic coefficient(75). We identified genes specific to the subtypes by computing the basis-specificity. The score for gene *i* was defined as:

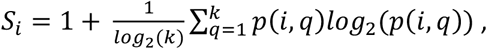

Where *k* is the factorization rank, and 𝑝(𝑖, 𝑞) is the probability that gene *i* contributes to cluster *q, i.e.* 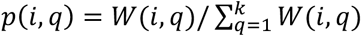.

The TCGA-LUAD expression subtype was identified as previously described(26). Briefly, the TMM normalized data was the median centered. The 506-gene signature for LUAD was applied by calculating the Pearson’s correlation of each sample to the centroids of the three TCGA-LUAD expression subtypes, respectively. The samples were assigned to the subtype with the highest correlation coefficient.

### Differential expression and gene set enrichment analyses

Differential expression analysis was performed using DESeq2 (RRID:SCR_015687) (76) to compare between one subtype versus all the other NS-LUAD tumors and repeated for each of the three subtypes. Genes with FDR < 0.05 and 50% change in expression level constituted the differentially expressed gene sets and were analyzed by IPA (RRID:SCR_008653) (24) to identify the enriched pathways. GSEA (RRID:SCR_003199) (25) was applied to test the gene sets associated with each subtype using the MSigDB (RRID:SCR_016863) hallmark gene sets(77) and an additional 155 curated gene sets associated with lung development(15). To exclude the possibility that the differential gene expression was the consequence of different tumor purity, we inferred the tumor purity from RNA-seq data using the R package ESTIMATE (RRID:SCR_026090) (78). The canonical marker genes were analyzed across three subtypes in samples with high tumor purity, defined as the purity estimate greater than 0.8.

### Inference of cell type compositions and histological lineages

For major cell types and alveolar epithelial cells, the gene signatures for deconvolution were obtained from a previous scRNA-seq analysis of lung cancer(41). The R package CAMTHC(79) was used to infer cell type abundances because of its high accuracy and robustness in a benchwork study of RNA-seq deconvolution methods(80). The deconvolution results of CAMTHC allows comparison across cell types.

For fibroblast cells, the gene signatures for deconvolution were obtained from a scRNA-seq study of stromal cells in lung cancer(36). The R package Bisque (RRID:SCR_005564) (42) was used to infer cell type abundances for its high accuracy. Of notes, this method estimates relative differences in abundances across samples that cannot be compared across cell types.

Cell lineage scores of LUAD, LUSC and NET subtypes were inferred from the RNA-seq data as described previously(33). Cancer histological type-specific genes were identified from scRNA-seq data(33). The score of each predicted histological type component in each LUAD was defined based on the total expression of genes specific to that histological type divided by the total expression of histological type-specific genes for all three subtypes. The mix-lineage score is defined as:

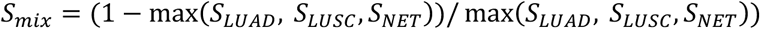

### Analysis of genomic driver events

504 RNA-seq tumor samples had WGS data available from the same subject. SNVs, SCNAs and global genomic features, including WGD and kataegis, were obtained from previous Sherlock-Lung publications(5,81,82). Fisher’s exact test was used to assess the association between genomic features and expression subtypes.

### Analysis of pathology image data

The diagnosis and morphological features of grade differentiation were reviewed by seven pathologists using H&E from the Sherlock*-Lung* FFPE tumor samples. Each tumor was reviewed by two pathologists independently. If there was not a consensus between the two pathologists, a third pathologist provided the final reading. In addition, six cytologic and immunophenotypic features were analyzed, including lymphoid aggregates, tumor spread through air spaces, clear cell feature, morular features, mucinous features and signet ring features. To ensure both sensitivity and specificity of detection, the presence of features was defined in two ways: (1) ‘OR’ logic: either of the two pathologists reported presence, and (2) ‘AND’ logic: both pathologists reported presence. The proportions of histologic components estimated by two pathologists were considered as consistent if the discrepancies were less than 20% and the mean values of the two estimates were used for the association analysis.

### Integrative analysis of cell compositions, expression subtypes and histologic components

In samples with consistent estimates of lepidic components, we evaluated the effect of the lepidic components on the association between cell compositions and expression subtypes. We fitted the generalized linear regression models (GLM) predicting the proportions of fibroblasts using the proportions of lepidic components and expression subtypes. P-values for individual subtype were assessed.

### Gene fusion analysis

Fusions were detected from RNA-Seq data using SFyNCS v0.15(83) and STAR-Fusion (RRID:SCR_025853) (84). For SFyNCS, the “duplication_like_and_inversion_like_distance” parameter was set to 300,000 so that duplication-like and inversion-like fusions with distance less than 300kb were filtered out. Default values were used for all other parameters. Fusions from 73 TCGA samples included in this study were obtained from the previous study(83). Additionally, fusions in three other TCGA samples (TCGA-44-2665, TCGA-50-5066, and TCGA-62-A46U) were detected using SFyNCS with the same parameters stated above. Fusion events detected by either software were included in the analysis and were annotated by annoFuse(85).

### Survival analysis

We investigated the associations of the three NS-LUAD expression subtypes with 10-year overall survival and additionally 5-year overall survival for the *chaotic* subtype using multivariate Cox proportional-hazards models adjusting for age, sex, and tumor stage (the stage was not included in the analysis of stage I subgroups). When evaluating the overall effect of the three subtypes, p-values were based on a likelihood ratio statistic, comparing models with and without the three subtypes, adjusting for the same covariates. When testing for the effect of a binary variable, p-values were based on a Wald statistic from Cox regression model. Survival analysis based on the 60-gene signature in the Sherlock data and the GIS data followed the same approach.

### Prediction of ICB response

The Sherlock data set was normalized by TPM after batch correction. RNA-seq data from 498 tumors with paired normal samples was normalized based on the paired normal samples. TIDE analysis was performed in the normalized data using the TIDE web application (RRID:SCR_026350) (57) (http://tide.dfci.harvard.edu/). The CTL level was estimated as the average expression level of *CD8A, CD8B, GZMA, GZMB* and *PRF1*. When evaluating the effect of three subtype on predicting TIDE score, we fitted the GLM model using immune gene expression (i.e. *PD-L1(CD274), PD-1(PDCD1*) or both) with or without the three subtypes. R-squared values for GLM and p-values based on a likelihood ratio statistic were assessed.

### Identification of the 60-gene signature

To identify a minimum gene set for expression subtype prediction, we applied a centroid-based method, Classification of Nearest Centroid (ClaNC)(86). ClaNC ranks genes by standard *t*-statistics and selects *N* genes for each cluster (aka, a total of 3\**N* genes for the three subtypes) to build the classifier. To identify the optimal *N*, we assessed 1 to 30 genes for each cluster by cross-validation and chose *N=*20 per subtype and a total of 60 genes for the signature of the three subtypes (**Supplementary Fig. S17A**). To evaluate the performance of this approach, we used cross-validation in the Sherlock data by randomly selecting 70% of samples as the training set to build the 60-gene signature. The predicted subtype was tested for error rate in the training set and the remaining 30% samples (testing set). The error rates for training and testing sets were reported based on 100 repeats of cross validation (**Supplementary Fig. S17B**). The final model was built using all Sherlock data and was applied to the GIS set for external validation. Samples were classified to a subtype based on the nearest centroid.

The risk models were trained with Cox regression in the Sherlock cohort using nine sets of variables, including (A) tumor stage, (B) driver mutations in *TP53*, *EGFR* and *KRAS*, and *EML4*-*ALK* fusions, (C) NS-LUAD expression subtypes, (D) histologic grade, (E) tumor stage plus mutations/fusions, (F) expression subtype plus stage, (G) expression subtype plus mutations/fusions, (H) expression subtype plus histologic grade, and (I) expression subtype plus both stage and mutations/fusions. For the metrics of expression subtypes, we used the distance to centroids. Specifically, for each subtype, we calculated the centroids of the 60 genes across all samples in this subtype in the training set. Then for each sample, we calculated the Euclidean distance to the centroids of the three subtypes, respectively. The coefficients of the Cox models were then applied to the GIS data to predict the risk scores. The C-indexes were calculated in the training and testing data sets, separately.

## Data Availability

The RNA-seq data have been deposited in SRA through dbGaP (RRID:SCR_002709) under the accession number phs003955.v1.p1. The major bioinformatic codes the NMF clustering, subtype prediction and survival analysis can be found at GitHub (RRID:SCR_002630) https://github.com/vivianzhao919/Sherlock-Lung.

## Authors’ Contributions

Conceptualization, MTL, WZ; Methodology, WZ, JSh, TZ, SA, XH, LY, MTL; Formal Analysis, WZ, XH, XZ, YY, MSah; Management: PHH; Pathology work: CL, KM, LMS, MKB, PJ, RH, SL, WDT, SY; Resources, DCC, KL, MPW, MMa, JL, BŚ, AM, OS, DZ, VJ, DM, SM, SO, MSav, MK, VG, PB, OA, YB, ESE, MBS, PH, LM, SSY, CC, CAH, IC, GL, JMS, BEG, NR, QL, MTL; Data Curation, PHoa, MMi, WZ; Writing – Original Draft, WZ, MTL; Writing – Review & Editing, All authors; Visualization: WZ, TZ, MTL; Supervision, MTL.

## Acknowledgements

This research was supported by the Intramural Research Program of the National Institute of Health (NIH) (project ZIACP101231 to MTL), and by the Anne Wojcicki Foundation (Grant Number: LC009). The contributions of the NIH author(s) were made as part of their official duties as NIH federal employees, are in compliance with agency policy requirements, and are considered Works of the United States Government. However, the findings and conclusions presented in this paper are those of the author(s) and do not necessarily reflect the views of the NIH or the U.S. Department of Health and Human Services. Where authors are identified as personnel of the International Agency for Research on Cancer/World Health Organization, the authors alone are responsible for the views expressed in this article and they do not necessarily represent the decisions, policy or views of the International Agency for Research on Cancer /World Health Organization. We want to acknowledge the patients and the INCLIVA Biobank (PT17/0015/0049) integrated in the Spanish National Biobanks Network and in the Valencian Biobanking Network for their collaboration. This study was supported by the Health and Medical Research Fund of Hong Kong SAR, HMRF 03142856. The related studies of Taiwan site were supported by grants from the Ministry of Health and Welfare, Taiwan DOH97-TD-G-111-026 (C.A.H.), DOH98-TD-G-111-015 (C.A.H.), DOH99-TD-G-111-028 (C.A.H.); DOH97-TD-G-111-029 (C.Y.C.), DOH98-TD-G-111-018 (C.Y.C.), DOH99-TD-G-111-015 (C.Y.C.), DOH97-TD-G-111-028(I.S.C.), DOH98-TD-G-111-017(I.S.C.), DOH99-TD-G-111-014(I.S.C.), and the Ministry of Science and Technology, Taiwan MOST109-2740-B-400-002 (C.A.H.), MOST110-2740-B-400-002 (C.A.H.), MOST111-2740-B-400-002 (C.A.H.). This work has been supported in part by the Tissue Core at the H. Lee Moffitt Cancer Center & Research Institute, a comprehensive cancer center designated by the National Cancer Institute and funded in part by a Moffitt Cancer Center Support Grant (no. P30-CA076292). And, in part, by NIH (NCI) grant # U01CA209414 to the Boston Lung Cancer Survival Study of the Dana-Farber/ Harvard Cancer Center (D.C.C.). The authors would like to thank the team at the IUCPQ site of the Quebec Respiratory Health Network Biobank of the FRQS for their valuable assistance, and would like to thank the staff at Harvard University, Yale University, Roswell Park Cancer Institute and Roswell PI, Instituto Nacional de Cancerologia, Nice University Hospital Centre (Nice UHC) - University Côte d’Azur and the Nice Biobank CRB, Toronto University Health Network, and Mayo Clinic for their assistance providing samples and corresponding clinical data. The computational analyses reported in this manuscript have utilized the NIH high-performance Biowulf Cluster. We thank the study participants and the staff at Westat Inc. for their assistance in collecting samples and corresponding clinical data. This study makes use of data generated by the Genome Institute of Singapore(22). This study was supported in part by the Memorial Sloan Kettering NCI core grant P30 CA008748.

## Supplementary Table Legends

**Supplementary Table S1.** Summary of RNA-seq-based studies for NS-LUAD.

**Supplementary Table S2.** Characteristics and clinical data of patients in the Sherlock cohort.

**Supplementary Table S3.** Summary of demographic and clinical characteristics of the Sherlock cohort.

**Supplementary Table S4.** NS-LUAD expression subtype-specific genes.

**Supplementary Table S5.** Gene signatures of hallmark genes and lung developmental pathways.

**Supplementary Table S6.** Histological type scores of the Sherlock cohort.

**Supplementary Table S7.** Fusion events detected in the Sherlock cohort.

**Supplementary Table S8.** Centroids of the 60-gene signature for the classification of NS-LUAD expression subtype.

**Supplementary Table S9.** NS-LUAD expression subtypes predicted by the 60-gene signature in the Sherlock cohort.

**Supplementary Table S10.** NS-LUAD expression subtypes predicted by the 60-gene signature in the GIS cohort.

## Supplementary Figure Legends

**Supplementary Fig. S1 Comparison of (A) age at diagnosis, (B) sex and (C) genetic ancestry of patients from the Sherlock-Lung and TCGA-LUAD studies in the Sherlock cohort.**

**Supplementary Fig. S2 Comparison of differentially expressed genes between subjects of East Asian (EAS) and European (EUR) ancestry.** The MA-plot shows differentially expressed genes in EUR relative to EAS NS-LUAD according to log ratio of EUR to EAS (y-axis) and average read counts (x-axis) of individual genes. Blue points indicate differentially expressed genes, defined as genes with FDR<0.05 and 50% change in expression level.

**Supplementary Fig. S3 Estimation of the number of clusters (factorization rank) for NMF clustering.** Consensus matrix for NMF rank 2 to 10.

**Supplementary Fig. S4 Pathway enrichment analysis of the Sherlock cohort. A,C,E,** Summary of IPA analysis of differentially expressed genes associated with the (**A**) *steady*, (**C**) *proliferative* and (**E**) *chaotic* subtypes, respectively. Orange and blue colors indicate positive and negative associations, respectively. **B,D,F,** Summary of GSEA analysis of gene sets associated with the (**B**) *steady*, (**D**) *proliferative* and (**F**) *chaotic* subtypes, respectively (FDR < 0.1). The bar plots show the normalized enrichment scores. FDR-adjusted p-values are indicated by colors. The pathway annotations are indicated in the left panels.

**Supplementary Fig. S5 GSEA enrichment plots of gene sets and expression levels of signature genes significantly associated with NS-LUAD expression subtypes. A,B,D,** Enrichment plots of top gene sets associated with *steady* related to (**A**) cell cycle, (**B**) stemness and (**D**) alveolar cells. The normalized enrichment scores (NES), p-values and FDRs are indicated in the plots. **c**, Violin plots depict the expression levels of the proliferation marker *MKI67*. (**E**) Enrichment plot of the top gene set associated with *proliferative* related to proliferation. **F-G**, Enrichment plots of top gene sets associated with *chaotic* related to (**F**) EMT, (**G**) fibroblasts. **H-I**, Violin plots depict expression levels of the mesenchymal markers (**H**) *CDH2* and (**I**) *TWIST1*. Mean values are indicated by the yellow lines in the violin plots. P-values from two-sided Mann-Whitney U-test are shown in corresponding figures. See Supplementary Table S5 for annotation of gene signatures.

**Supplementary Fig. S6 Comparison of the canonical proliferation and mesenchymal marker genes across NS-LUAD expression subtypes in tumor samples of high purity.** Violin plots depict the expression levels of (**A**) the proliferation marker *MKI67*, and the mesenchymal markers (**B**) *CDH2* and (**C**) *TWIST1* in tumor samples with estimated purity of greater than 0.8. Mean values are indicated by the yellow lines in the violin plots. P-values from two-sided Mann-Whitney U-test are shown in corresponding figures.

**Supplementary Fig. S7 S-LUAD expression subtypes in the Sherlock cohort.** (**A**) Clustering of 506-gene centroid-based classifier of S-LUAD expression subtypes. The top panel shows data set, sex, genetic ancestry, S-LUAD subtypes, and NS-LUAD expression subtypes. (**B**) Proportions of S-LUAD expression subtypes across three NS-LUAD expression subtypes.

**Supplementary Fig. S8 Comparison of clear cell features across the NS-LUAD expression subtypes.** P-values from chi-squared test are shown.

**Supplementary Fig. S9 Comparison of histological cell lineages across tumor stages in the Sherlock cohort.** Violin plots depict the LUAD histology scores across tumor stages in (**A**) all tumors and (**B**) within *steady*, *proliferative* and *chaotic* subtypes, respectively. Mean values are indicated by the yellow lines. P-values from ordinal test are shown.

**Supplementary Fig. S10 Comparison of the proportions of fibroblasts and the proportions of lepidic component.** NS-LUAD expression subtypes are indicated by colors. P-value from Pearson correlation test is shown.

**Supplementary Fig. S11 Comparison of proportions of (A) epithelial cells, (B) fibroblasts, (C) AT2 cells and (D) basal cells across NS-LUAD expression subtypes stratified by tumor stages.** Mean values are indicated by the yellow lines. P-values from two-sided Mann-Whitney U-test are shown.

**Supplementary Fig. S12 Comparison of expression levels of cell type signature genes across NS-LUAD expression subtypes.** Violin plots depict signature genes for (**A-B**) fibroblasts (*COL1A1*, *PDGFRA*), (**C**) epithelial cells (*EPCAM*), (**D-E**) AT2 cells (*SFTPB*, *SFTPC*), (**F-G**) basal cells (*KRT17*, *FHL2*), (**H-I**) *COL10A1*+ and *COL4A1*+ cancer associated fibroblasts (*COL10A1*, *COL4A1*) and (**J**) non-malignant fibroblasts (*VEGFD*). Mean values are indicated by the yellow lines. P-values from two-sided Mann-Whitney U-test are shown.

**Supplementary Fig. S13 Summary of genomic alterations in the NS-LUAD expression subtypes. A-C**, Summary of cancer driver genes and additional genomic alterations associated with (**A**) *steady*, (**B**) *proliferative* and (**C**) *chaotic* subtypes. The volcano plots show significant associations according to p-values (y-axis) and odds ratio based on Fisher’s exact test (x-axis). The colors and sizes of the points indicate types and frequencies of the alterations, respectively. The green and red lines indicate FDR = 0.1 and p = 0.05, respectively.

**Supplementary Fig. S14 Comparison of genomic alterations in NS-LUAD expression subtypes.** Bar plots depict the proportions of (**A**) *TP53* driver mutations, (**B**) whole genome doubling (WGD) status, (**C**) *ALK* fusions, and (**D**) *EGFR* driver mutations in *steady*, *proliferative* and *chaotic* subtypes.

**Supplementary Fig. S15 Fusion events detected in the Sherlock cohort. A-B**, Proportions of (**A**) interchromosome and intrachromosome fusions and (**B**) fusions supported by structural variants (SV) across NS-LUAD expression subtypes. (**C**) Summary of types of genes involved in fusions. Circle sizes indicate the numbers of genes. **D-E**, (**D**) Numbers of protein-coding gene fusions per sample and (**E**) numbers of in-frame gene fusions per sample across NS-LUAD expression subtypes. Mean values are indicated by the yellow lines. P-values from two-sided Mann-Whitney U-test are shown.

**Supplementary Fig. S16 Association between NS-LUAD expression subtypes and predicted response to immune checkpoint blockade (ICB) therapy.** (**A**) Comparison of TIDE scores predicting response to ICB across NS-LUAD expression subtypes. **B-D**, Comparison of (**B**) predicted cytotoxic T lymphocyte (CTL) levels, (**C**) myeloid-derived suppressor cells (MDSC) and (**D**) cancer associated fibroblast (CAF) abundances across NS-LUAD expression subtypes. Mean values are indicated by the yellow lines. P-values from two-sided Mann-Whitney U-test are shown. **E-F**, Correlation between TIDE scores and (**E**) *PD-L1*(*CD274*) and (**F**) *PD-1*(*PDCD1*) expression levels. NS-LUAD expression subtypes are indicated by colors and shapes. Based on the known proportion of ICB response in non-small cell lung cancers^61^, the bottom 20% (lowest quintile) of the TIDE scores are indicated by the dashed lines. Marginal distribution maps of gene levels and TIDE scores stratified by expression subtypes are displayed at the top and side panels of the plots. (**G**) Comparison of R-squared values from models predicting TIDE scores using gene expression of *PD-L1*, *PD-1*, and *PD-L1* plus *PD-1* with and without the information on transcriptomic-based subtypes.

**Supplementary Fig. S17 Identification of the 60-gene signature for classification of NS-LUAD expression subtypes.** (**A**) Estimation of the minimum gene sets per NS-LUAD expression subtype by cross validation in the Sherlock cohort. 1 to 30 genes per subtype were assessed. Box plots represent the numbers of genes per NS-LUAD expression subtype (x-axis) and the error rates defined by cross-validation (y-axis). (**B**) Distribution of the error rates for predicting NS-LUAD expression subtypes using a 60-gene signature by cross validation in the Sherlock cohort. The error rates in the training and testing sets are indicated by colors. The median error rates are indicated by dashed lines.

**Supplementary Fig. S18 Association between NS-LUAD expression subtypes predicted by the 60-gene signature and overall survival in the Sherlock cohort. A-D**, Kaplan-Meier survival curves for overall survival stratified by the predicted expression subtypes in (**A**) all patients, (**B**) stage I patients, (**C**) patients classified as TRU subtype^26^, and (**D**) *TP53*-wt patients. P-values and hazard ratios (HR) are calculated using Cox proportional hazards models adjusting for age, sex, and tumor stage (the analysis restricted to tumor stage I did not include tumor stage in the model).

## References

1. Cufari ME, Proli C, De Sousa P, Raubenheimer H, Al Sahaf M, Chavan H, et al. Increasing frequency of non-smoking lung cancer: Presentation of patients with early disease to a tertiary institution in the UK. Eur J Cancer 2017;84:55–9 doi 10.1016/j.ejca.2017.06.031.

2. Pelosof L, Ahn C, Gao A, Horn L, Madrigales A, Cox J, et al. Proportion of Never-Smoker Non-Small Cell Lung Cancer Patients at Three Diverse Institutions. J Natl Cancer Inst 2017;109(7) doi 10.1093/jnci/djw295.

3. Sung H, Ferlay J, Siegel RL, Laversanne M, Soerjomataram I, Jemal A, Bray F. Global Cancer Statistics 2020: GLOBOCAN Estimates of Incidence and Mortality Worldwide for 36 Cancers in 185 Countries. CA Cancer J Clin 2021;71(3):209–49 doi 10.3322/caac.21660.

4. Sun S, Schiller JH, Gazdar AF. Lung cancer in never smokers--a different disease. Nat Rev Cancer 2007;7(10):778–90 doi 10.1038/nrc2190.

5. Zhang T, Joubert P, Ansari-Pour N, Zhao W, Hoang PH, Lokanga R, et al. Genomic and evolutionary classification of lung cancer in never smokers. Nat Genet 2021;53(9):1348–59 doi 10.1038/s41588-021-00920-0.

6. LoPiccolo J, Gusev A, Christiani DC, Janne PA. Lung cancer in patients who have never smoked - an emerging disease. Nat Rev Clin Oncol 2024;21(2):121–46 doi 10.1038/s41571-023-00844-0.

7. Nam AS, Chaligne R, Landau DA. Integrating genetic and non-genetic determinants of cancer evolution by single-cell multi-omics. Nat Rev Genet 2021;22(1):3–18 doi 10.1038/s41576-020-0265-5.

8. Yoshida K, Gowers KHC, Lee-Six H, Chandrasekharan DP, Coorens T, Maughan EF, et al. Tobacco smoking and somatic mutations in human bronchial epithelium. Nature 2020;578(7794):266–72 doi 10.1038/s41586-020-1961-1.

9. Martincorena I, Roshan A, Gerstung M, Ellis P, Van Loo P, McLaren S, et al. Tumor evolution. High burden and pervasive positive selection of somatic mutations in normal human skin. Science 2015;348(6237):880–6 doi 10.1126/science.aaa6806.

10. Shaffer SM, Dunagin MC, Torborg SR, Torre EA, Emert B, Krepler C, et al. Rare cell variability and drug-induced reprogramming as a mode of cancer drug resistance. Nature 2017;546(7658):431–5 doi 10.1038/nature22794.

11. Emert BL, Cote CJ, Torre EA, Dardani IP, Jiang CL, Jain N, et al. Variability within rare cell states enables multiple paths toward drug resistance. Nat Biotechnol 2021;39(7):865–76 doi 10.1038/s41587-021-00837-3.

12. Wang Z, Li Z, Zhou K, Wang C, Jiang L, Zhang L, et al. Deciphering cell lineage specification of human lung adenocarcinoma with single-cell RNA sequencing. Nat Commun 2021;12(1):6500 doi 10.1038/s41467-021-26770-2.

13. Marjanovic ND, Hofree M, Chan JE, Canner D, Wu K, Trakala M, et al. Emergence of a High-Plasticity Cell State during Lung Cancer Evolution. Cancer Cell 2020;38(2):229–46 e13 doi 10.1016/j.ccell.2020.06.012.

14. Juul NH, Yoon JK, Martinez MC, Rishi N, Kazadaeva YI, Morri M, et al. KRAS(G12D) drives lepidic adenocarcinoma through stem-cell reprogramming. Nature 2023;619(7971):860–7 doi 10.1038/s41586-023-06324-w.

15. Laughney AM, Hu J, Campbell NR, Bakhoum SF, Setty M, Lavallee VP, et al. Regenerative lineages and immune-mediated pruning in lung cancer metastasis. Nat Med 2020;26(2):259–69 doi 10.1038/s41591-019-0750-6.

16. Zhao W, Zhu B, Hutchinson A, Pesatori AC, Consonni D, Caporaso NE, et al. Clinical Implications of Inter-and Intratumor Heterogeneity of Immune Cell Markers in Lung Cancer. J Natl Cancer Inst 2022;114(2):280–9 doi 10.1093/jnci/djab157.

17. Cancer Genome Atlas Research Network. Comprehensive molecular profiling of lung adenocarcinoma. Nature 2014;511(7511):543–50 doi 10.1038/nature13385.

18. Group PTC, Calabrese C, Davidson NR, Demircioglu D, Fonseca NA, He Y, et al. Genomic basis for RNA alterations in cancer. Nature 2020;578(7793):129–36 doi 10.1038/s41586-020-1970-0.

19. Martinez-Ruiz C, Black JRM, Puttick C, Hill MS, Demeulemeester J, Larose Cadieux E, et al. Genomic-transcriptomic evolution in lung cancer and metastasis. Nature 2023;616(7957):543–52 doi 10.1038/s41586-023-05706-4.

20. Rosenthal R, Cadieux EL, Salgado R, Bakir MA, Moore DA, Hiley CT, et al. Neoantigen-directed immune escape in lung cancer evolution. Nature 2019;567(7749):479–85 doi 10.1038/s41586-019-1032-7.

21. Shi J, Hua X, Zhu B, Ravichandran S, Wang M, Nguyen C, et al. Somatic Genomics and Clinical Features of Lung Adenocarcinoma: A Retrospective Study. PLoS Med 2016;13(12):e1002162 doi 10.1371/journal.pmed.1002162.

22. Chen J, Yang H, Teo ASM, Amer LB, Sherbaf FG, Tan CQ, et al. Genomic landscape of lung adenocarcinoma in East Asians. Nat Genet 2020;52(2):177–86 doi 10.1038/s41588-019-0569-6.

23. Devarajan K. Nonnegative matrix factorization: an analytical and interpretive tool in computational biology. PLoS Comput Biol 2008;4(7):e1000029 doi 10.1371/journal.pcbi.1000029.

24. QIAGEN Inc. Available from: https://www.qiagenbioinformatics.com/products/ingenuity-pathway-analysis

25. Subramanian A, Tamayo P, Mootha VK, Mukherjee S, Ebert BL, Gillette MA, et al. Gene set enrichment analysis: a knowledge-based approach for interpreting genome-wide expression profiles. Proc Natl Acad Sci U S A 2005;102(43):15545–50 doi 10.1073/pnas.0506580102.

26. Wilkerson MD, Yin X, Walter V, Zhao N, Cabanski CR, Hayward MC, et al. Differential pathogenesis of lung adenocarcinoma subtypes involving sequence mutations, copy number, chromosomal instability, and methylation. PLoS One 2012;7(5):e36530 doi 10.1371/journal.pone.0036530.

27. Bhattacharjee A, Richards WG, Staunton J, Li C, Monti S, Vasa P, et al. Classification of human lung carcinomas by mRNA expression profiling reveals distinct adenocarcinoma subclasses. Proc Natl Acad Sci U S A 2001;98(24):13790–5 doi 10.1073/pnas.191502998.

28. Takeuchi T, Tomida S, Yatabe Y, Kosaka T, Osada H, Yanagisawa K, et al. Expression profile-defined classification of lung adenocarcinoma shows close relationship with underlying major genetic changes and clinicopathologic behaviors. J Clin Oncol 2006;24(11):1679–88 doi 10.1200/JCO.2005.03.8224.

29. Hayes DN, Monti S, Parmigiani G, Gilks CB, Naoki K, Bhattacharjee A, et al. Gene expression profiling reveals reproducible human lung adenocarcinoma subtypes in multiple independent patient cohorts. J Clin Oncol 2006;24(31):5079–90 doi 10.1200/JCO.2005.05.1748.

30. Tsuta K, Ishii G, Yoh K, Nitadori J, Hasebe T, Nishiwaki Y, et al. Primary lung carcinoma with signet-ring cell carcinoma components: clinicopathological analysis of 39 cases. Am J Surg Pathol 2004;28(7):868–74 doi 10.1097/00000478-200407000-00004.

31. Ou SH, Ziogas A, Zell JA. Primary signet-ring carcinoma (SRC) of the lung: a population-based epidemiologic study of 262 cases with comparison to adenocarcinoma of the lung. J Thorac Oncol 2010;5(4):420–7 doi 10.1097/JTO.0b013e3181ce3b93.

32. Iwasaki T, Ohta M, Lefor AT, Kawahara K. Signet-ring cell carcinoma component in primary lung adenocarcinoma: potential prognostic factor. Histopathology 2008;52(5):639–40 doi 10.1111/j.1365-2559.2008.02987.x.

33. Li Q, Wang R, Yang Z, Li W, Yang J, Wang Z, et al. Molecular profiling of human non-small cell lung cancer by single-cell RNA-seq. Genome Med 2022;14(1):87 doi 10.1186/s13073-022-01089-9.

34. Xing X, Yang F, Huang Q, Guo H, Li J, Qiu M, et al. Decoding the multicellular ecosystem of lung adenocarcinoma manifested as pulmonary subsolid nodules by single-cell RNA sequencing. Sci Adv 2021;7(5) doi 10.1126/sciadv.abd9738.

35. Goveia J, Rohlenova K, Taverna F, Treps L, Conradi LC, Pircher A, et al. An Integrated Gene Expression Landscape Profiling Approach to Identify Lung Tumor Endothelial Cell Heterogeneity and Angiogenic Candidates. Cancer Cell 2020;37(1):21–36 e13 doi 10.1016/j.ccell.2019.12.001.

36. Lambrechts D, Wauters E, Boeckx B, Aibar S, Nittner D, Burton O, et al. Phenotype molding of stromal cells in the lung tumor microenvironment. Nat Med 2018;24(8):1277–89 doi 10.1038/s41591-018-0096-5.

37. Leader AM, Grout JA, Maier BB, Nabet BY, Park MD, Tabachnikova A, et al. Single-cell analysis of human non-small cell lung cancer lesions refines tumor classification and patient stratification. Cancer Cell 2021;39(12):1594–609 e12 doi 10.1016/j.ccell.2021.10.009.

38. Wu F, Fan J, He Y, Xiong A, Yu J, Li Y, et al. Single-cell profiling of tumor heterogeneity and the microenvironment in advanced non-small cell lung cancer. Nat Commun 2021;12(1):2540 doi 10.1038/s41467-021-22801-0.

39. Kim N, Kim HK, Lee K, Hong Y, Cho JH, Choi JW, et al. Single-cell RNA sequencing demonstrates the molecular and cellular reprogramming of metastatic lung adenocarcinoma. Nat Commun 2020;11(1):2285 doi 10.1038/s41467-020-16164-1.

40. Zilionis R, Engblom C, Pfirschke C, Savova V, Zemmour D, Saatcioglu HD, et al. Single-Cell Transcriptomics of Human and Mouse Lung Cancers Reveals Conserved Myeloid Populations across Individuals and Species. Immunity 2019;50(5):1317–34 e10 doi 10.1016/j.immuni.2019.03.009.

41. Maynard A, McCoach CE, Rotow JK, Harris L, Haderk F, Kerr DL, et al. Therapy-Induced Evolution of Human Lung Cancer Revealed by Single-Cell RNA Sequencing. Cell 2020;182(5):1232–51 e22 doi 10.1016/j.cell.2020.07.017.

42. Jew B, Alvarez M, Rahmani E, Miao Z, Ko A, Garske KM, et al. Accurate estimation of cell composition in bulk expression through robust integration of single-cell information. Nat Commun 2020;11(1):1971 doi 10.1038/s41467-020-15816-6.

43. Yang P, Kulig K, Boland JM, Erickson-Johnson MR, Oliveira AM, Wampfler J, et al. Worse disease-free survival in never-smokers with ALK+ lung adenocarcinoma. J Thorac Oncol 2012;7(1):90–7 doi 10.1097/JTO.0b013e31823c5c32.

44. Parikh K, Dimou A, Leventakos K, Mansfield AS, Shanshal M, Wan Y, et al. Impact of EML4-ALK Variants and Co-Occurring TP53 Mutations on Duration of First-Line ALK Tyrosine Kinase Inhibitor Treatment and Overall Survival in ALK Fusion-Positive NSCLC: Real-World Outcomes From the GuardantINFORM database. J Thorac Oncol 2024;19(11):1539–49 doi 10.1016/j.jtho.2024.07.009.

45. Aisner DL, Sholl LM, Berry LD, Rossi MR, Chen H, Fujimoto J, et al. The Impact of Smoking and TP53 Mutations in Lung Adenocarcinoma Patients with Targetable Mutations-The Lung Cancer Mutation Consortium (LCMC2). Clin Cancer Res 2018;24(5):1038–47 doi 10.1158/1078-0432.CCR-17-2289.

46. Mortusewicz O, Fouquerel E, Ame JC, Leonhardt H, Schreiber V. PARG is recruited to DNA damage sites through poly(ADP-ribose)-and PCNA-dependent mechanisms. Nucleic Acids Res 2011;39(12):5045–56 doi 10.1093/nar/gkr099.

47. Piscuoglio S, Ng CKY, Geyer FC, Burke KA, Cowell CF, Martelotto LG, et al. Genomic and transcriptomic heterogeneity in metaplastic carcinomas of the breast. NPJ Breast Cancer 2017;3:48 doi 10.1038/s41523-017-0048-0.

48. Ahrendt SA, Hu Y, Buta M, McDermott MP, Benoit N, Yang SC, et al. p53 mutations and survival in stage I non-small-cell lung cancer: results of a prospective study. J Natl Cancer Inst 2003;95(13):961–70 doi 10.1093/jnci/95.13.961.

49. Steels E, Paesmans M, Berghmans T, Branle F, Lemaitre F, Mascaux C, et al. Role of p53 as a prognostic factor for survival in lung cancer: a systematic review of the literature with a meta-analysis. Eur Respir J 2001;18(4):705–19 doi 10.1183/09031936.01.00062201.

50. Liam CK, Yew CY, Pang YK, Wong CK, Poh ME, Tan JL, et al. Common driver mutations and programmed death-ligand 1 expression in advanced non-small cell lung cancer in smokers and never smokers. BMC Cancer 2023;23(1):659 doi 10.1186/s12885-023-11156-y.

51. Rizvi NA, Mazieres J, Planchard D, Stinchcombe TE, Dy GK, Antonia SJ, et al. Activity and safety of nivolumab, an anti-PD-1 immune checkpoint inhibitor, for patients with advanced, refractory squamous non-small-cell lung cancer (CheckMate 063): a phase 2, single-arm trial. Lancet Oncol 2015;16(3):257–65 doi 10.1016/S1470-2045(15)70054-9.

52. Taube JM, Klein A, Brahmer JR, Xu H, Pan X, Kim JH, et al. Association of PD-1, PD-1 ligands, and other features of the tumor immune microenvironment with response to anti-PD-1 therapy. Clin Cancer Res 2014;20(19):5064–74 doi 10.1158/1078-0432.CCR-13-3271.

53. Brahmer JR, Drake CG, Wollner I, Powderly JD, Picus J, Sharfman WH, et al. Phase I study of single-agent anti-programmed death-1 (MDX-1106) in refractory solid tumors: safety, clinical activity, pharmacodynamics, and immunologic correlates. J Clin Oncol 2010;28(19):3167–75 doi 10.1200/JCO.2009.26.7609.

54. Yarchoan M, Hopkins A, Jaffee EM. Tumor Mutational Burden and Response Rate to PD-1 Inhibition. N Engl J Med 2017;377(25):2500–1 doi 10.1056/NEJMc1713444.

55. Rizvi NA, Hellmann MD, Snyder A, Kvistborg P, Makarov V, Havel JJ, et al. Cancer immunology. Mutational landscape determines sensitivity to PD-1 blockade in non-small cell lung cancer. Science 2015;348(6230):124–8 doi 10.1126/science.aaa1348.

56. Ricciuti B, Wang X, Alessi JV, Rizvi H, Mahadevan NR, Li YY, et al. Association of High Tumor Mutation Burden in Non-Small Cell Lung Cancers With Increased Immune Infiltration and Improved Clinical Outcomes of PD-L1 Blockade Across PD-L1 Expression Levels. JAMA Oncol 2022;8(8):1160–8 doi 10.1001/jamaoncol.2022.1981.

57. Jiang P, Gu S, Pan D, Fu J, Sahu A, Hu X, et al. Signatures of T cell dysfunction and exclusion predict cancer immunotherapy response. Nat Med 2018;24(10):1550–8 doi 10.1038/s41591-018-0136-1.

58. Gajewski TF, Schreiber H, Fu YX. Innate and adaptive immune cells in the tumor microenvironment. Nat Immunol 2013;14(10):1014–22 doi 10.1038/ni.2703.

59. Joyce JA, Fearon DT. T cell exclusion, immune privilege, and the tumor microenvironment. Science 2015;348(6230):74–80 doi 10.1126/science.aaa6204.

60. Topalian SL, Taube JM, Anders RA, Pardoll DM. Mechanism-driven biomarkers to guide immune checkpoint blockade in cancer therapy. Nat Rev Cancer 2016;16(5):275–87 doi 10.1038/nrc.2016.36.

61. Xue L, Bi G, Zhan C, Zhang Y, Yuan Y, Fan H. Development and Validation of a 12-Gene Immune Relevant Prognostic Signature for Lung Adenocarcinoma Through Machine Learning Strategies. Front Oncol 2020;10:835 doi 10.3389/fonc.2020.00835.

62. Wang X, Yao S, Xiao Z, Gong J, Liu Z, Han B, Zhang Z. Development and validation of a survival model for lung adenocarcinoma based on autophagy-associated genes. J Transl Med 2020;18(1):149 doi 10.1186/s12967-020-02321-z.

63. Wang Z, Embaye KS, Yang Q, Qin L, Zhang C, Liu L, et al. Establishment and validation of a prognostic signature for lung adenocarcinoma based on metabolism-related genes. Cancer Cell Int 2021;21(1):219 doi 10.1186/s12935-021-01915-x.

64. Lai YH, Chen WN, Hsu TC, Lin C, Tsao Y, Wu S. Overall survival prediction of non-small cell lung cancer by integrating microarray and clinical data with deep learning. Sci Rep 2020;10(1):4679 doi 10.1038/s41598-020-61588-w.

65. Huang Z, Shi M, Zhou H, Wang J, Zhang HJ, Shi J. Prognostic signature of lung adenocarcinoma based on stem cell-related genes. Sci Rep 2021;11(1):1687 doi 10.1038/s41598-020-80453-4.

66. Dobin A, Davis CA, Schlesinger F, Drenkow J, Zaleski C, Jha S, et al. STAR: ultrafast universal RNA-seq aligner. Bioinformatics 2013;29(1):15–21 doi 10.1093/bioinformatics/bts635.

67. Anders S, Pyl PT, Huber W. HTSeq--a Python framework to work with high-throughput sequencing data. Bioinformatics 2015;31(2):166–9 doi 10.1093/bioinformatics/btu638.

68. Frankish A, Carbonell-Sala S, Diekhans M, Jungreis I, Loveland JE, Mudge JM, et al. GENCODE: reference annotation for the human and mouse genomes in 2023. Nucleic Acids Res 2023;51(D1):D942–D9 doi 10.1093/nar/gkac1071.

69. Andrews S. Available from: http://www.bioinformatics.babraham.ac.uk/projects/fastqc

70. Broad Institute. Available from: http://broadinstitute.github.io/picard

71. Grossman RL, Heath AP, Ferretti V, Varmus HE, Lowy DR, Kibbe WA, Staudt LM. Toward a Shared Vision for Cancer Genomic Data. N Engl J Med 2016;375(12):1109–12 doi 10.1056/NEJMp1607591.

72. Zhang Y, Parmigiani G, Johnson WE. ComBat-seq: batch effect adjustment for RNA-seq count data. NAR Genom Bioinform 2020;2(3):lqaa078 doi 10.1093/nargab/lqaa078.

73. Love MI, Huber W, Anders S. Moderated estimation of fold change and dispersion for RNA-seq data with DESeq2. Genome Biol 2014;15(12):550 doi 10.1186/s13059-014-0550-8.

74. Gaujoux R, Seoighe C. A flexible R package for nonnegative matrix factorization. BMC Bioinformatics 2010;11:367 doi 10.1186/1471-2105-11-367.

75. Brunet JP, Tamayo P, Golub TR, Mesirov JP. Metagenes and molecular pattern discovery using matrix factorization. Proc Natl Acad Sci U S A 2004;101(12):4164–9 doi 10.1073/pnas.0308531101.

76. Chen L. CAMTHC: Convex Analysis of Mixtures for Tissue Heterogeneity Characterization. R package version 1.2.0. 2019.

77. Liberzon A, Subramanian A, Pinchback R, Thorvaldsdottir H, Tamayo P, Mesirov JP. Molecular signatures database (MSigDB) 3.0. Bioinformatics 2011;27(12):1739–40 doi 10.1093/bioinformatics/btr260.

78. Yoshihara K, Shahmoradgoli M, Martinez E, Vegesna R, Kim H, Torres-Garcia W, et al. Inferring tumour purity and stromal and immune cell admixture from expression data. Nat Commun 2013;4:2612 doi 10.1038/ncomms3612.

79. Chen L. CAMTHC: convex analysis of mixtures for tissue heterogeneity characterization. 2019.

80. Jin H, Liu Z. A benchmark for RNA-seq deconvolution analysis under dynamic testing environments. Genome Biol 2021;22(1):102 doi 10.1186/s13059-021-02290-6.

81. Zhang T, Sang J, Hoang PH, Zhao W, Rosenbaum J, Johnson KE, et al. APOBEC shapes tumor evolution and age at onset of lung cancer in smokers. bioRxiv 2024 doi 10.1101/2024.04.02.587805.

82. Diaz-Gay M, Zhang T, Hoang PH, Khandekar A, Zhao W, Steele CD, et al. The mutagenic forces shaping the genomic landscape of lung cancer in never smokers. medRxiv 2024 doi 10.1101/2024.05.15.24307318.

83. Zhong X, Luan J, Yu A, Lee-Hassett A, Miao Y, Yang L. SFyNCS detects oncogenic fusions involving non-coding sequences in cancer. Nucleic Acids Res 2023;51(18):e96 doi 10.1093/nar/gkad705.

84. Haas BJ, Dobin A, Li B, Stransky N, Pochet N, Regev A. Accuracy assessment of fusion transcript detection via read-mapping and de novo fusion transcript assembly-based methods. Genome Biol 2019;20(1):213 doi 10.1186/s13059-019-1842-9.

85. Gaonkar KS, Marini F, Rathi KS, Jain P, Zhu Y, Chimicles NA, et al. annoFuse: an R Package to annotate, prioritize, and interactively explore putative oncogenic RNA fusions. BMC Bioinformatics 2020;21(1):577 doi 10.1186/s12859-020-03922-7.

86. Dabney AR. Classification of microarrays to nearest centroids. Bioinformatics 2005;21(22):4148–54 doi 10.1093/bioinformatics/bti681.

