## Supplementary Figures for "A prognostic signature for lung adenocarcinoma in people who have never smoked"

### Supplementary Fig. S1

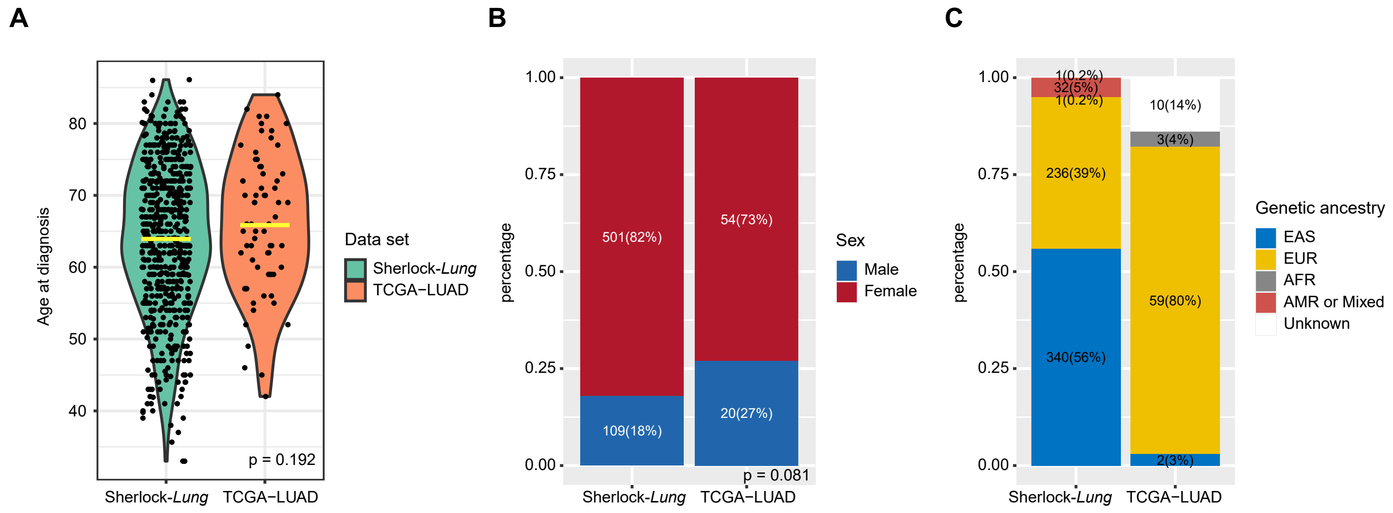


**Supplementary Fig. S1 Comparison of (A) age at diagnosis, (B) sex and (C) genetic ancestry of patients from the Sherlock-*Lung* and TCGA-LUAD studies in the Sherlock cohort.**

**Supplementary Fig. S2**

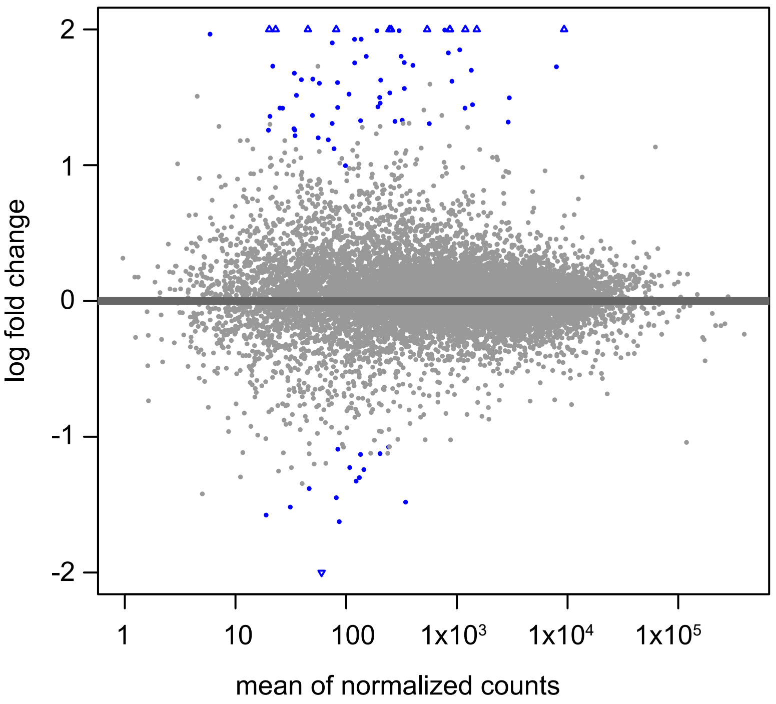


**Supplementary Fig. S2 Comparison of differentially expressed genes between subjects of East Asian (EAS) and European (EUR) ancestry.** The MA-plot shows differentially expressed genes in EUR relative to EAS NS-LUAD according to log ratio of EUR to EAS (y-axis) and average read counts (x-axis) of individual genes. Blue points indicate differentially expressed genes, defined as genes with FDR<0.05 and 50% change in expression level.

### Supplementary Fig. S3

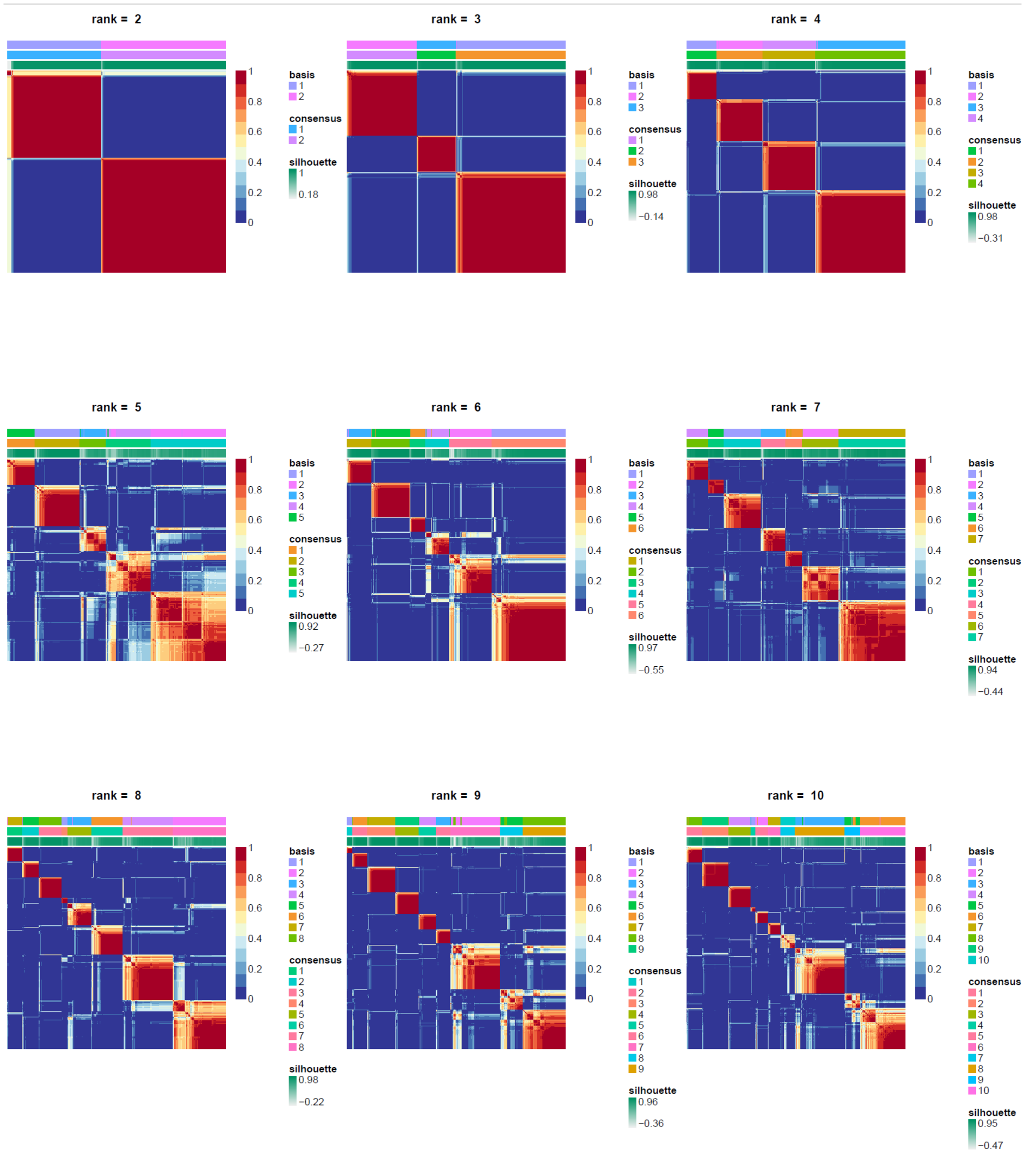


**Supplementary Fig. S3 Estimation of the number of clusters (factorization rank) for NMF clustering.** Consensus matrix for NMF rank 2 to 10.

### Supplementary Fig. S4

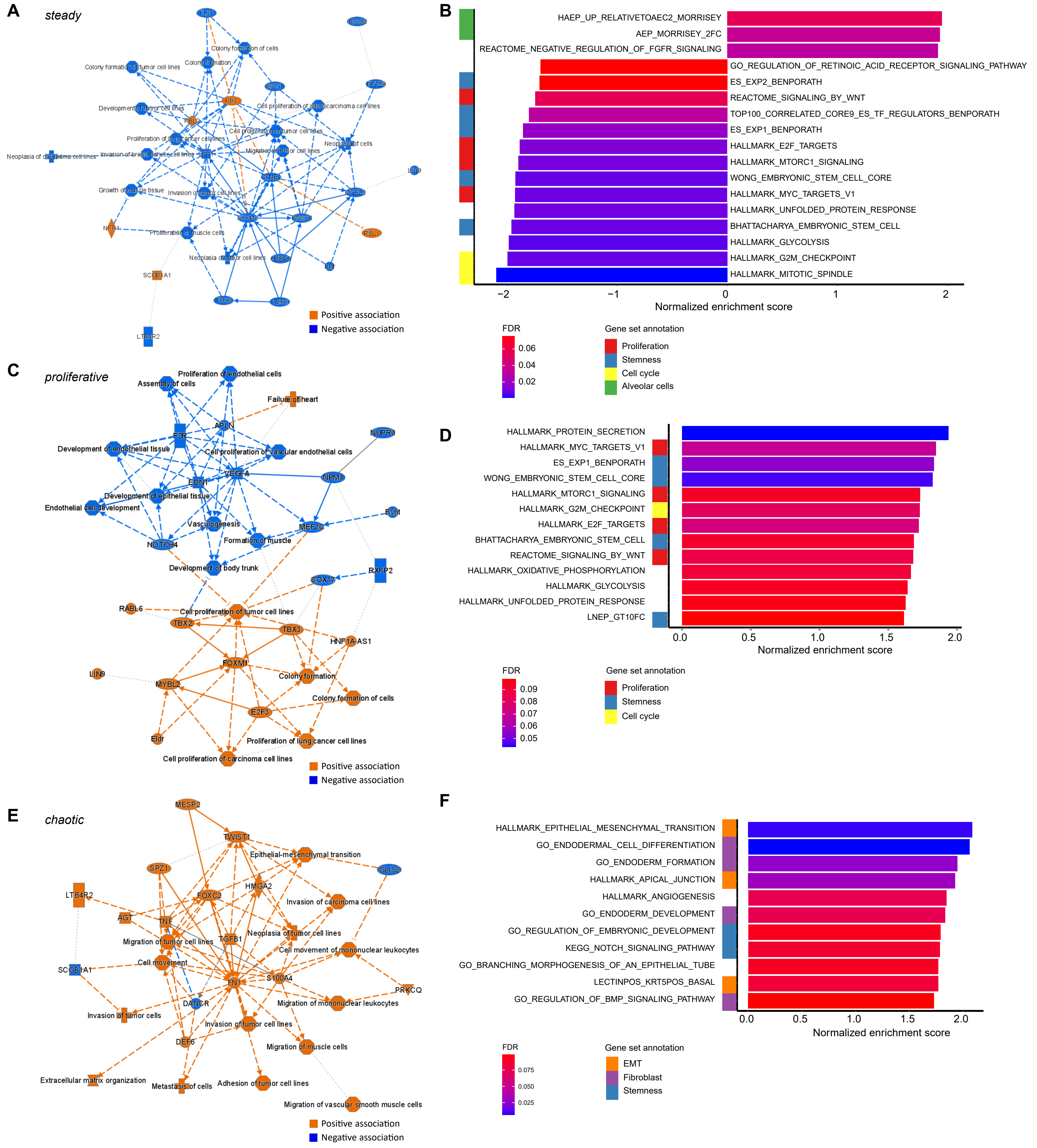


**Supplementary Fig. S4 Pathway enrichment analysis of the Sherlock cohort. A,C,E,** Summary of IPA analysis of differentially expressed genes associated with the (**A**) *steady*, (**C**) *proliferative* and (**E**) *chaotic* subtypes, respectively. Orange and blue colors indicate positive and negative associations, respectively. **B,D,F,** Summary of GSEA analysis of gene sets associated with the (**B**) *steady*, (**D**) *proliferative* and (**F**) *chaotic* subtypes, respectively (FDR < 0.1). The bar plots show the normalized enrichment scores. FDR-adjusted p-values are indicated by colors. The pathway annotations are indicated in the left panels.

#### Supplementary Fig. S5

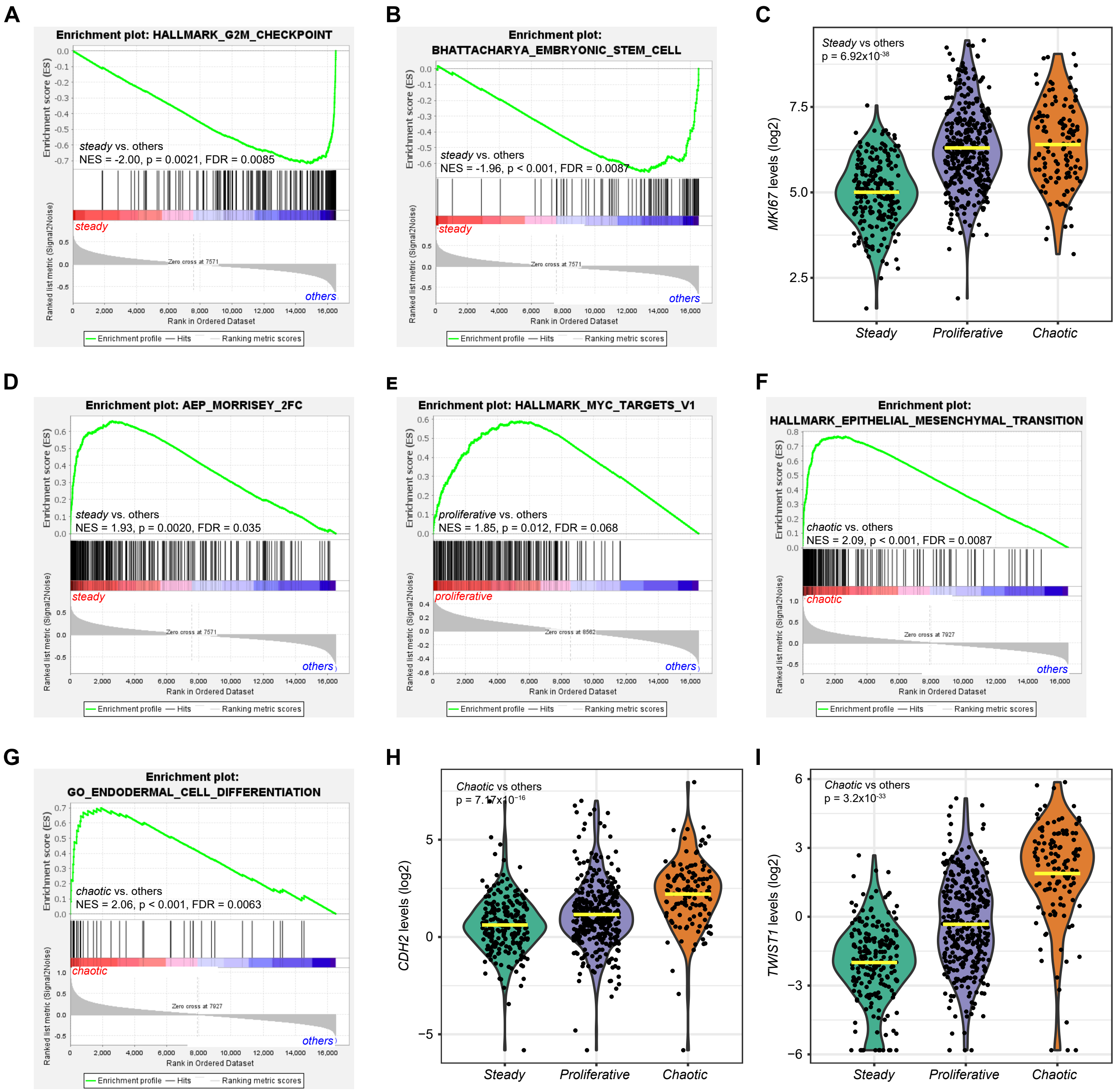


**Supplementary Fig. S5 GSEA enrichment plots of gene sets and expression levels of signature genes significantly associated with NS-LUAD expression subtypes. A,B,D**, Enrichment plots of top gene sets associated with *steady* related to (**A**) cell cycle, (**B**) stemness and (**D**) alveolar cells. The normalized enrichment scores (NES), p-values and FDRs are indicated in the plots. **C**, Violin plots depict the expression levels of the proliferation marker *MKI67*. (**E**) Enrichment plot of the top gene set associated with *proliferative* related to proliferation. **F-G**, Enrichment plots of top gene sets associated with *chaotic* related to (**F**) EMT, (**G**) fibroblasts. **H-I**, Violin plots depict expression levels of the mesenchymal markers (**H**) *CDH2* and (**I**) *TWIST1*. Mean values are indicated by the yellow lines in the violin plots. P-values from two-sided Mann-Whitney U-test are shown in corresponding figures. See Supplementary Table S5 for annotation of gene signatures.

#### Supplementary Fig. S6

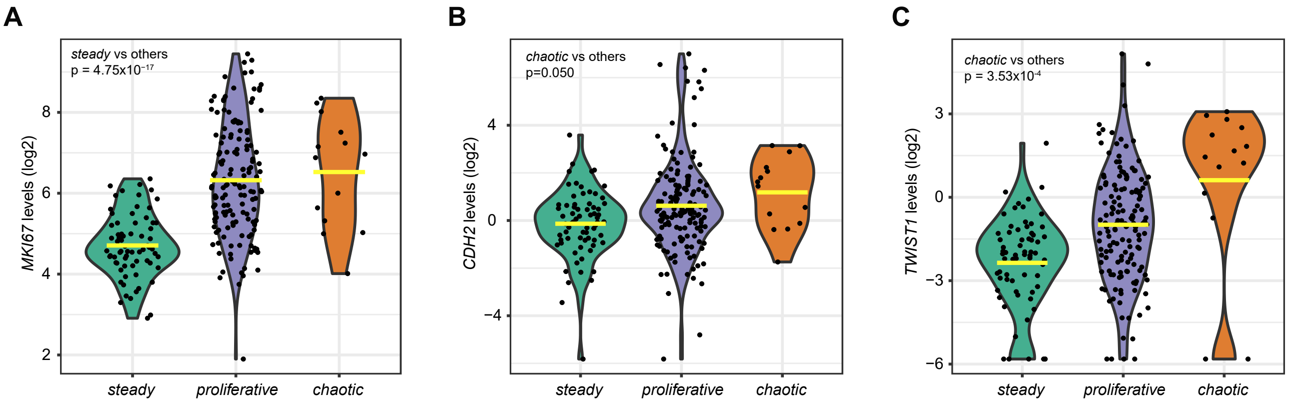


**Supplementary Fig. S6 Comparison of the canonical proliferation and mesenchymal marker genes across NS-LUAD expression subtypes in tumor samples of high purity.** Violin plots depict the expression levels of (**A**) the proliferation marker *MKI67*, and the mesenchymal markers (**B**) *CDH2* and (**C**) *TWIST1* in tumor samples with estimated purity of greater than 0.8. Mean values are indicated by the yellow lines in the violin plots. P-values from two-sided Mann-Whitney U-test are shown in corresponding figures.

#### Supplementary Fig. S7

A

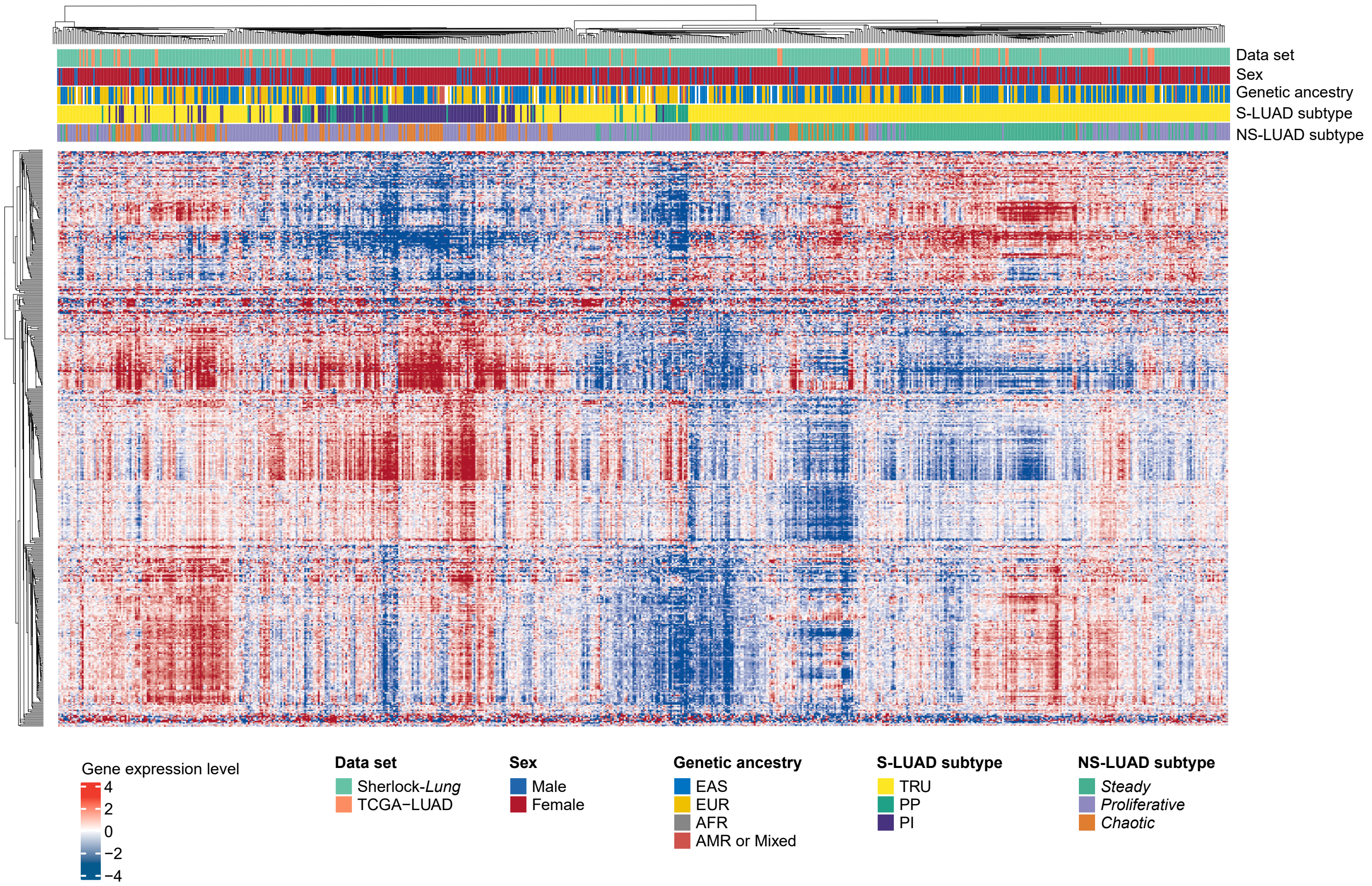


B

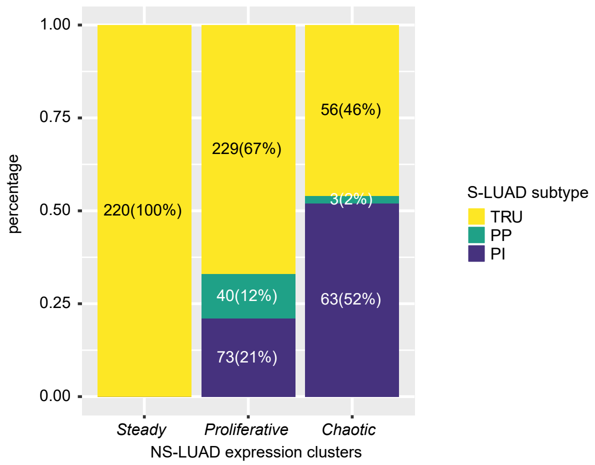


**Supplementary Fig. S7 S-LUAD expression subtypes in the Sherlock cohort.** (**A**) Clustering of 506-gene centroid-based classifier of S-LUAD expression subtypes. The top panel shows data set, sex, genetic ancestry, S-LUAD subtypes, and NS-LUAD expression subtypes. (**B**) Proportions of S-LUAD expression subtypes across three NS-LUAD expression subtypes.

#### Supplementary Fig. S8

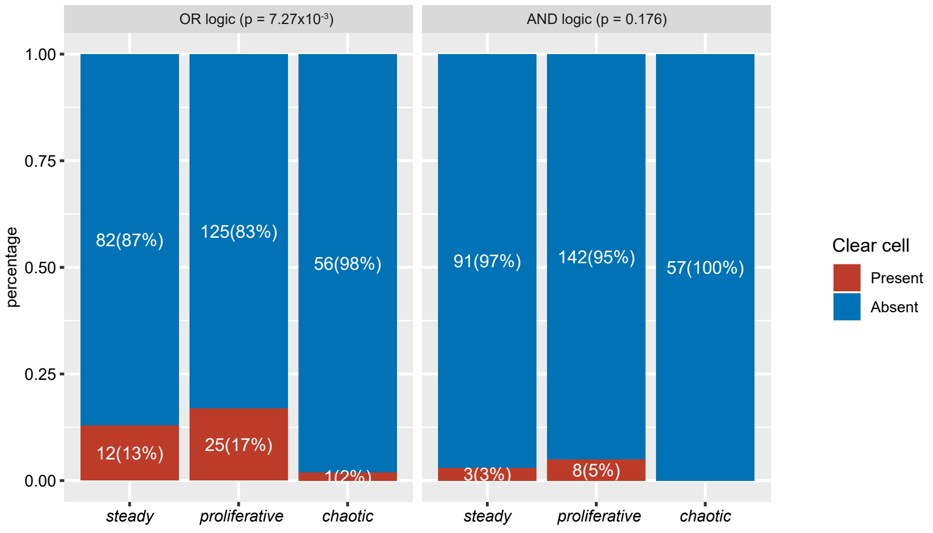


**Supplementary Fig. S8 Comparison of clear cell features across the NS-LUAD expression subtypes.** P-values from chi-squared test are shown.

#### Supplementary Fig. S9

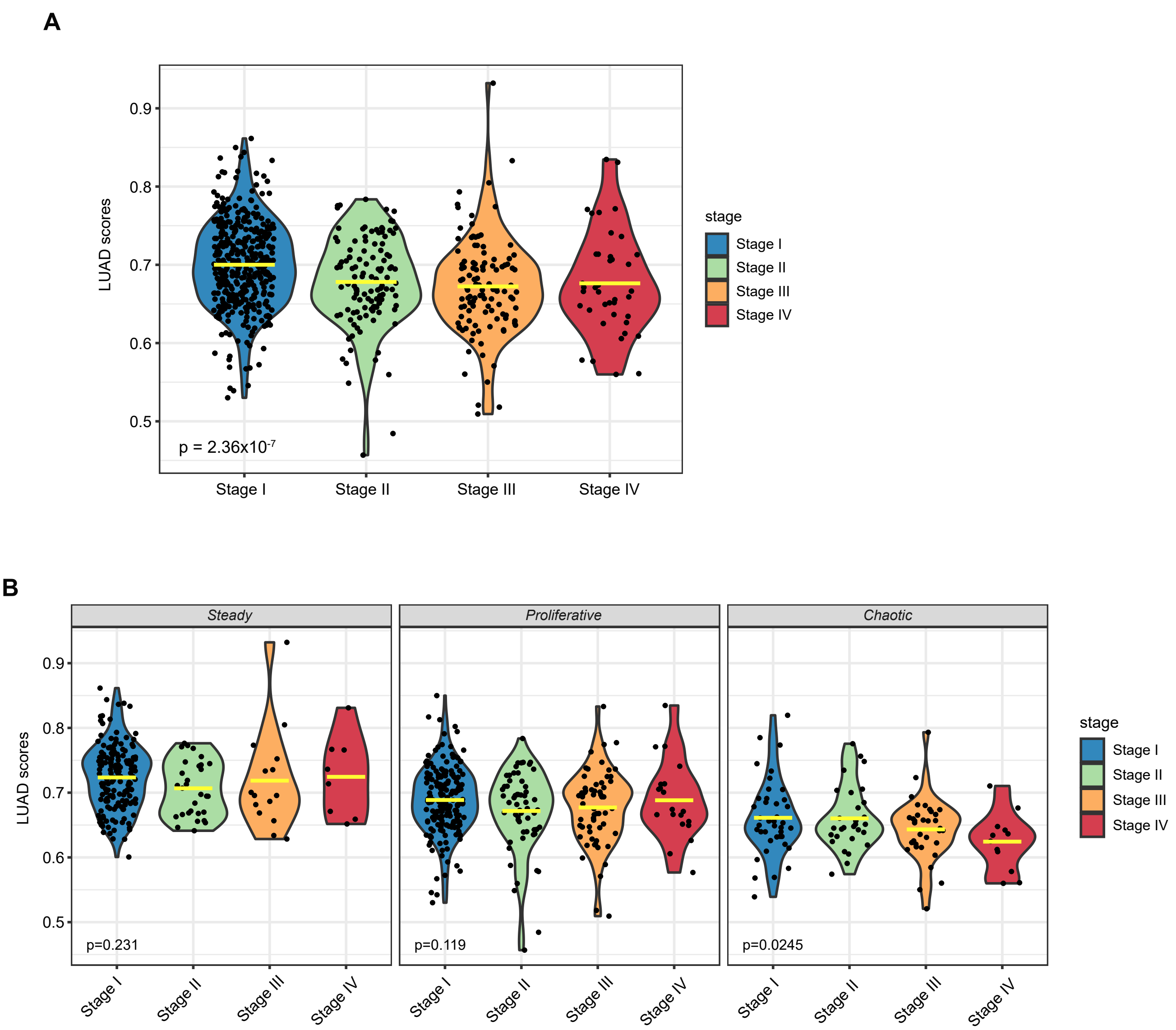


**Supplementary Fig. S9 Comparison of histological cell lineages across tumor stages in the Sherlock cohort.** Violin plots depict the LUAD histology scores across tumor stages in (**A**) all tumors and (**B**) within *steady*, *proliferative* and *chaotic* subtypes, respectively. Mean values are indicated by the yellow lines. P-values from ordinal test are shown.

#### Supplementary Fig. S10

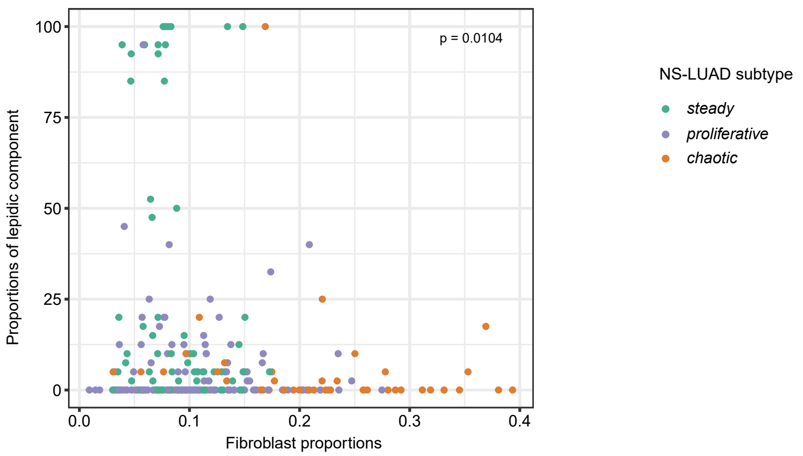


**Supplementary Fig. S10 Comparison of the proportions of fibroblasts and the proportions of lepidic component.** NS-LUAD expression subtypes are indicated by colors. P-value from Pearson correlation test is shown.

#### Supplementary Fig. S11

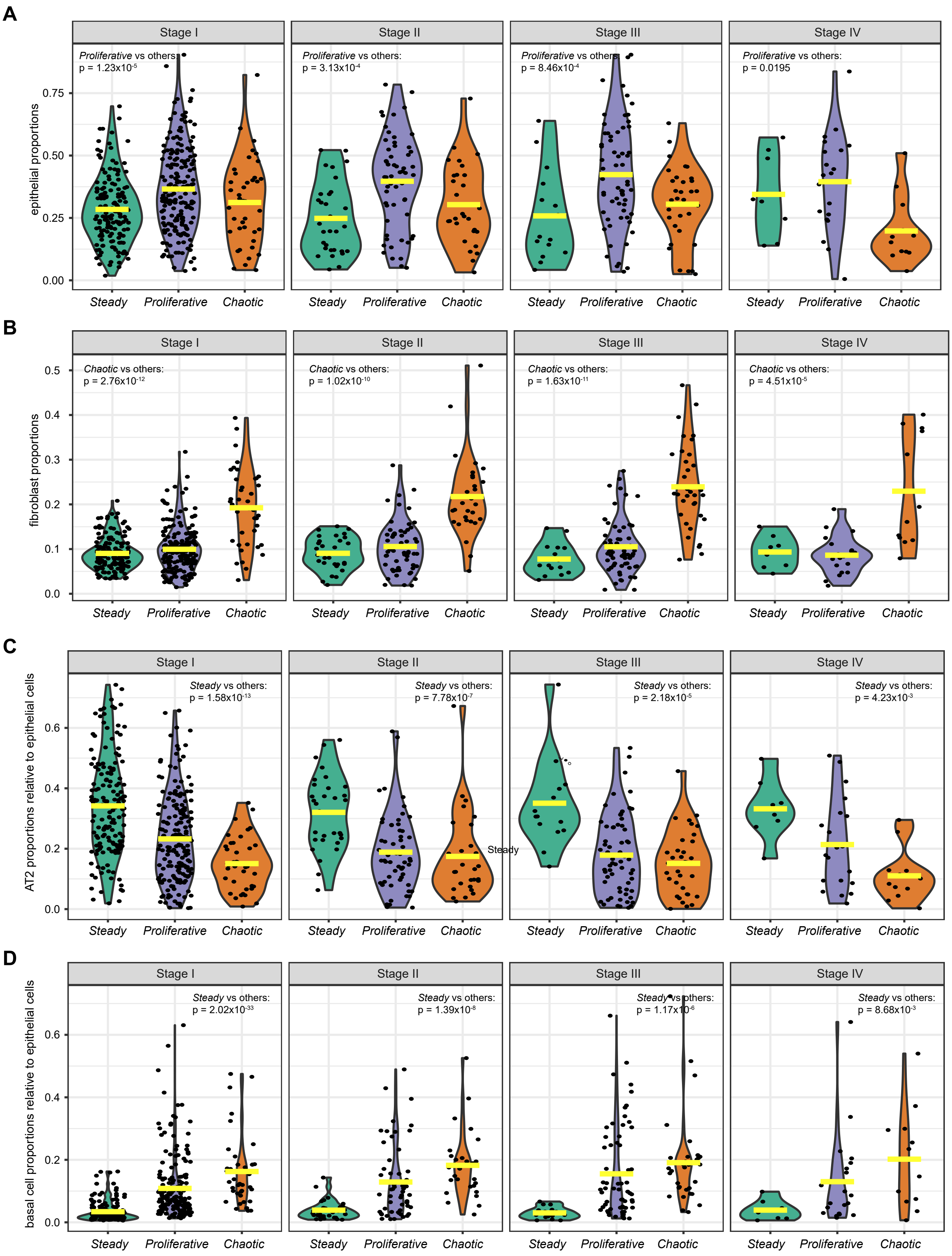


**Supplementary Fig. S11 Comparison of proportions of (A) epithelial cells, (B) fibroblasts, (C) AT2 cells and (D) basal cells across NS-LUAD expression subtypes stratified by tumor stages.** Mean values are indicated by the yellow lines. P-values from two-sided Mann-Whitney U-test are shown.

#### Supplementary Fig. S12

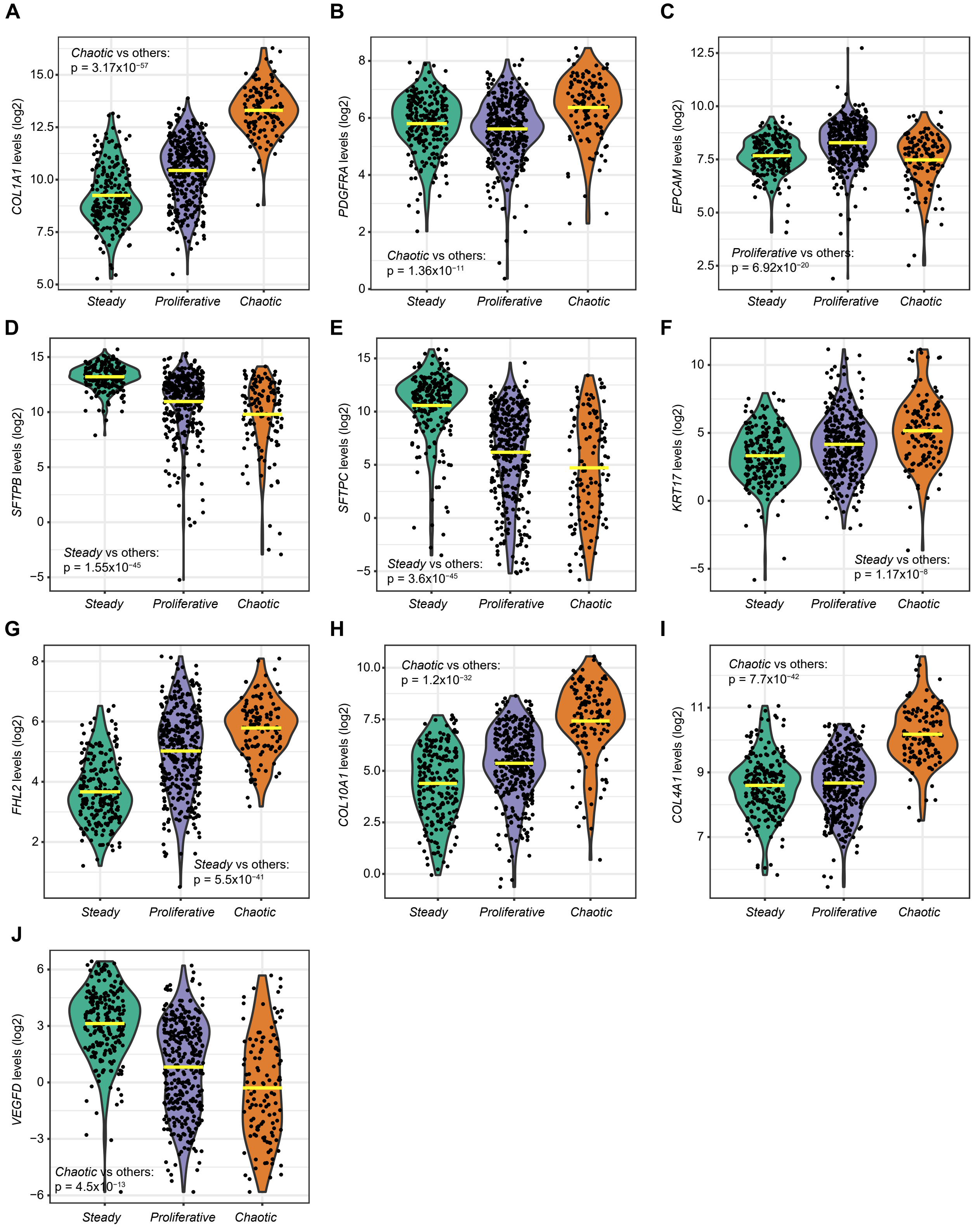


**Supplementary Fig. S12 Comparison of expression levels of cell type signature genes across NS-LUAD expression subtypes.** Violin plots depict signature genes for (**A-B**) fibroblasts (*COL1A1*, *PDGFRA*), (**C**) epithelial cells (*EPCAM*), (**D-E**) AT2 cells (*SFTPB*, *SFTPC*), (**F-G**) basal cells (*KRT17*, *FHL2*), (**H-I**) *COL10A1*$^{+}$ and *COL4A1*$^{+}$ cancer associated fibroblasts (*COL10A1*, *COL4A1*) and (**J**) non-malignant fibroblasts (*VEGFD*). Mean values are indicated by the yellow lines. P-values from two-sided Mann-Whitney U-test are shown.

**Supplementary Fig. S13**

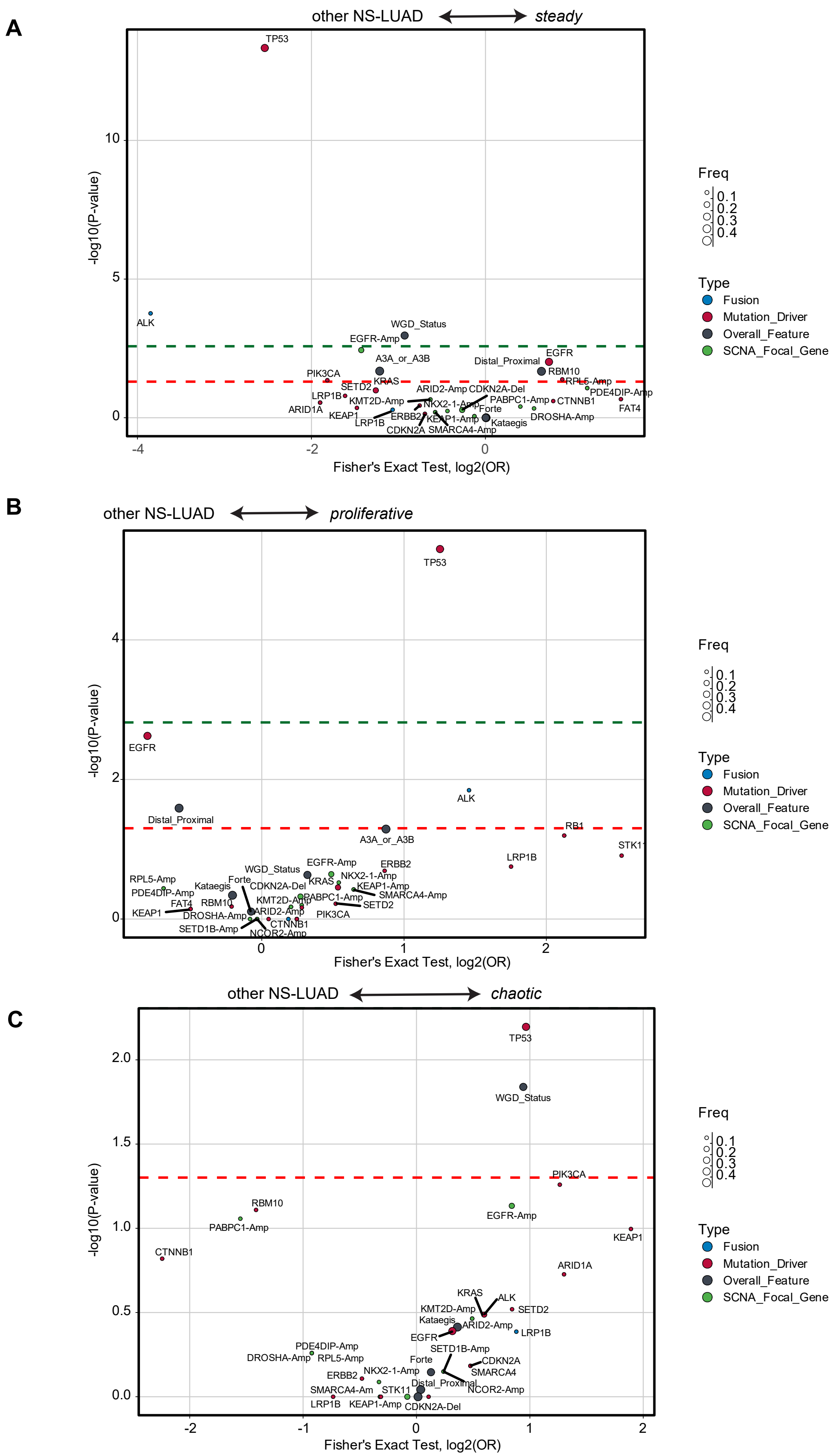


**Supplementary Fig. S13 Summary of genomic alterations in the NS-LUAD expression subtypes. A-C**, Summary of cancer driver genes and additional genomic alterations associated with (**A**) *steady*, (**B**) *proliferative* and (**C**) *chaotic* subtypes. The volcano plots show significant associations according to p-values (y-axis) and odds ratio based on Fisher's exact test (x-axis). The colors and sizes of the points indicate types and frequencies of the alterations, respectively. The green and red lines indicate FDR = 0.1 and p = 0.05, respectively.

#### Supplementary Fig. S14

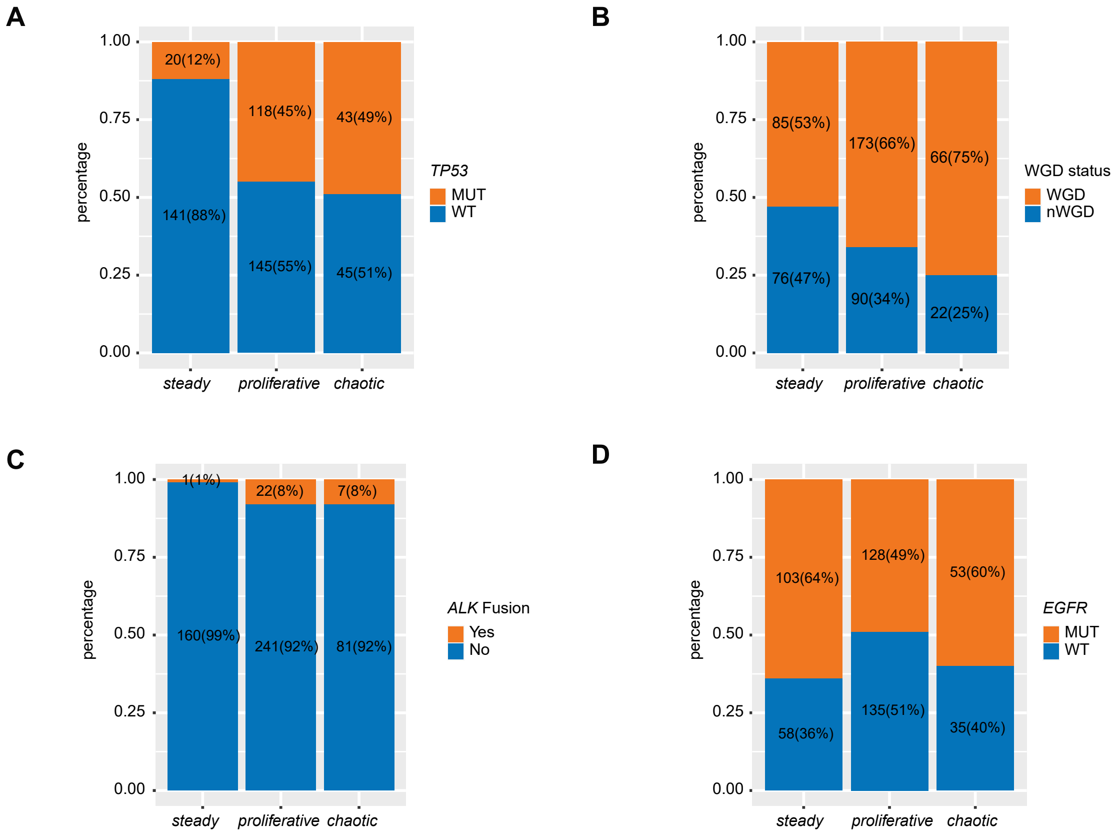


**Supplementary Fig. S14 Comparison of genomic alterations in NS-LUAD expression subtypes.** Bar plots depict the proportions of (**A**) *TP53* driver mutations, (**B**) whole genome doubling (WGD) status, (**C**) *ALK* fusions, and (**D**) *EGFR* driver mutations in *steady*, *proliferative* and *chaotic* subtypes.

### Supplementary Fig. S15

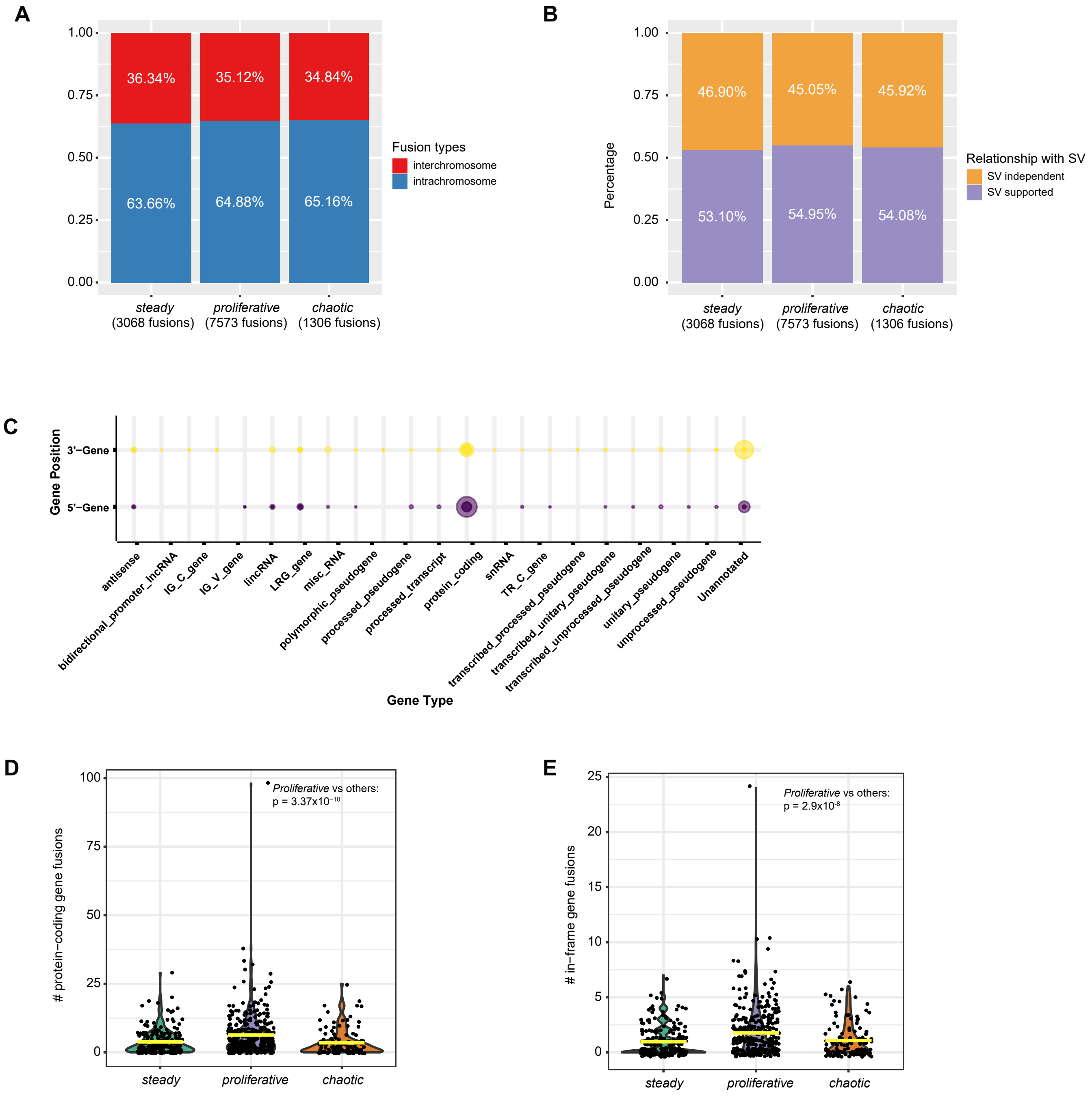


**Supplementary Fig. S15 Fusion events detected in the Sherlock cohort. A-B**, Proportions of (**A**) interchromosome and intrachromosome fusions and (**B**) fusions supported by structural variants (SV) across NS-LUAD expression subtypes. (**C**) Summary of types of genes involved in fusions. Circle sizes indicate the numbers of genes. **D-E**, (**D**) Numbers of protein-coding gene fusions per sample and (**E**) numbers of in-frame gene fusions per sample across NS-LUAD expression subtypes. Mean values are indicated by the yellow lines. P-values from two-sided Mann-Whitney U-test are shown.

### Supplementary Fig. S16

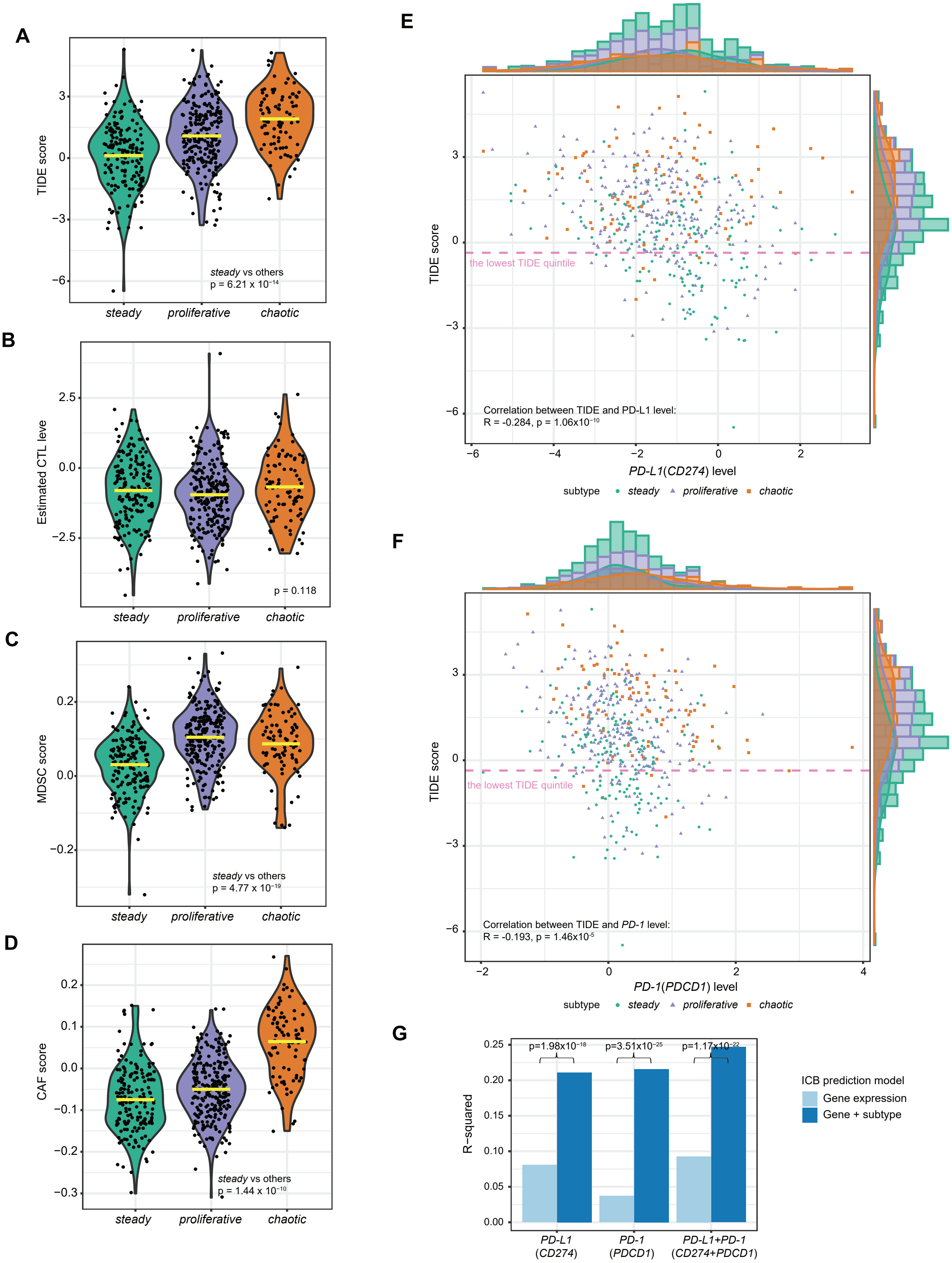

**Supplementary Fig. S16 Association between NS-LUAD expression subtypes and predicted response to immune checkpoint blockade (ICB) therapy.** (**A**) Comparison of TIDE scores predicting response to ICB across NS-LUAD expression subtypes. **B-D**, Comparison of (**B**) predicted cytotoxic T lymphocyte (CTL) levels, (**C**) myeloid-derived suppressor cells (MDSC) and (**D**) cancer associated fibroblast (CAF) abundances across NS-LUAD expression subtypes. Mean values are indicated by the yellow lines. P-values from two-sided Mann-Whitney U-test are shown. **E-F**, Correlation between TIDE scores and (**E**) *PD-L1*(*CD274*) and (**F**) *PD-1*(*PDCD1*) expression levels. NS-LUAD expression subtypes are indicated by colors and shapes. Based on the known proportion of ICB response in non-small cell lung cancers[61], the bottom 20% (lowest quintile) of the TIDE scores are indicated by the dashed lines. Marginal distribution maps of gene levels and TIDE scores stratified by expression subtypes are displayed at the top and side panels of the plots. (**G**) Comparison of R-squared values from models predicting TIDE scores using gene expression of *PD-L1*, *PD-1*, and *PD-L1* plus *PD-1* with and without the information on transcriptomic-based subtypes.

#### Supplementary Fig. S17

**A**

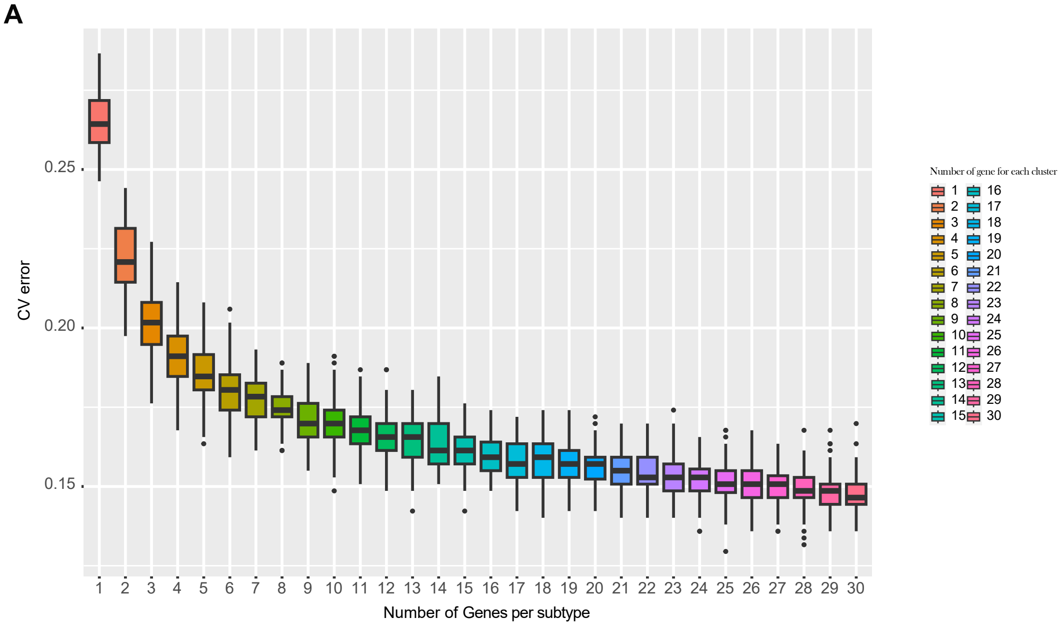


**B**

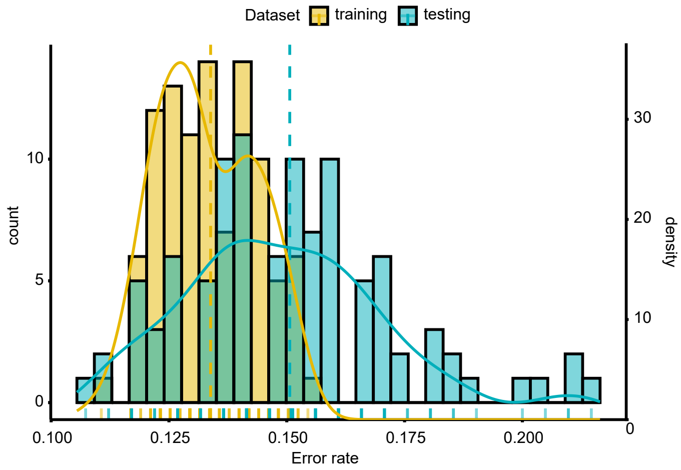


**Supplementary Fig. S17 Identification of the 60-gene signature for classification of NS-LUAD expression subtypes.** (**A**) Estimation of the minimum gene sets per NS-LUAD expression subtype by cross validation in the Sherlock cohort. 1 to 30 genes per subtype were assessed. Box plots represent the numbers of genes per NS-LUAD expression subtype (x-axis) and the error rates defined by cross-validation (y-axis). (**B**) Distribution of the error rates for predicting NS-LUAD expression subtypes using a 60-gene signature by cross validation in the Sherlock cohort. The error rates in the training and testing sets are indicated by colors. The median error rates are indicated by dashed lines.

#### Supplementary Fig. S18

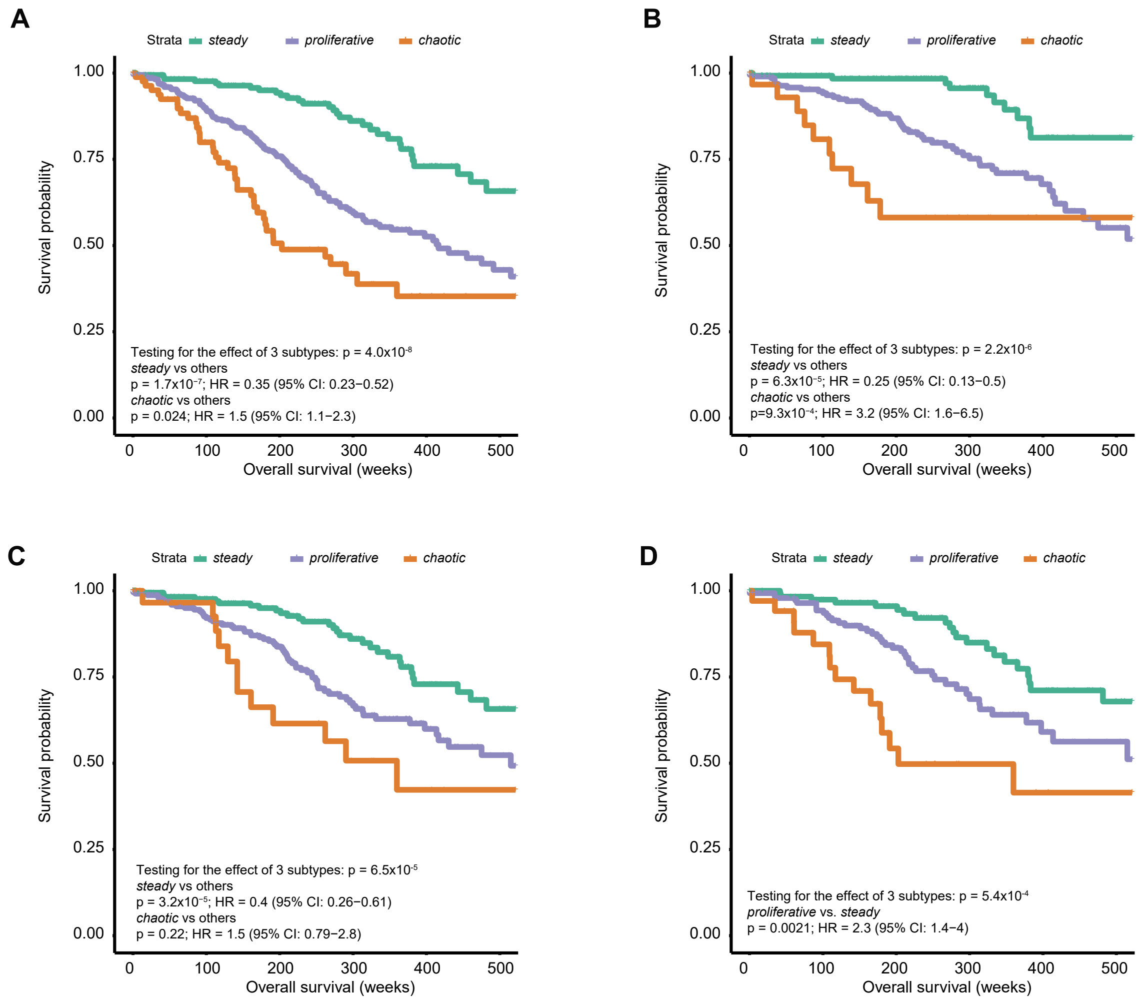


**Supplementary Fig. S18 Association between NS-LUAD expression subtypes predicted by the 60-gene signature and overall survival in the Sherlock cohort. A-D**, Kaplan-Meier survival curves for overall survival stratified by the predicted expression subtypes in (**A**) all patients, (**B**) stage **I** patients, (**C**) patients classified as TRU subtype[26], and (**D**) *TP53*-wt patients. P-values and hazard ratios (HR) are calculated using Cox proportional hazards models adjusting for age, sex, and tumor stage (the analysis restricted to tumor stage I did not include tumor stage in the model).
